# Genomic analyses provide insights into genetic architecture of three-way crossbred pigs

**DOI:** 10.1101/491753

**Authors:** Yu Lin, Qianzi Tang, Yan Li, Mengnan He, Long Jin, Jideng Ma, Xun Wang, Keren Long, Zhiqing Huang, Xuewei Li, Yiren Gu, Mingzhou Li

## Abstract

**Background:** Crossbreeding is effective for improving performance in poultry and livestock, which is mainly attributed to heterosis. For pork production, a classic three-way crossbreeding system of Duroc × (Landrace × Yorkshire) (DLY) is widely used to produce terminal crossbred pigs with stable and prominent performance. Nonetheless, studies on the transmission of genetic information and gene expression pattern of DLY have been limited.

**Findings:** We analyzed population-scale SNPs based on 30 individuals from these three purebreds and identified 529.93 K SNPs of breed-of-origin of alleles. We also applied whole-genome sequencing of ten individuals from a DLY pig family as well as transcriptome of four representative tissues (adipose, skeletal muscle, heart, and liver) for six DLY individuals. Based on above, we identified a large number of high-confidence ASE genes, among which four ASE genes (*KMO*, *PLIN4*, *POPDC3* and *UGT1A6*) were found to be shared over all DLY individuals.

**Conclusion:** We suggest DLY is a more effective strategy of three-way crossbreeding among these three purebreds from genetic aspect. We suppose the numerous breed-of-origin of alleles have close association with improved performance of crossbred individuals. ASE may also play important roles on DLY three-way crossbreeding system. Our findings are valuable for understanding the transmission of genetic information and the gene expression in DLY three-way crossbreeding and may be used to guide breeding and production of pigs in the future.

## Background

Crossbreeding for animals (especially agricultural livestock and poultry) is a classic and effective method to utilize heterosis, a phenomenon that progeny of diverse purebreds exhibits better performance than their parents [1,2,3]. Using crossbreeding, alleles that are specific to different purebreds will be inherited by the offspring and generate a large number of heterozygous single-nucleotide polymorphisms (SNPs). These would be suggested to improve crossbred performance, especially in terms of growth rate, reproductive performance, production performance (meat, egg, and milk), and disease resistance [3,4].

Pigs (*Sus scrofa*) were domesticated at least ~9,000 years ago and have been used as a major source of animal protein in the human diet; there are over 730 distinct pig breeds worldwide [5,6]. The worldwide distribution of pigs is dominated by six international transboundary commercial pig breeds originating in Europe, namely, Berkshire, Duroc, Hampshire, Landrace, Piétrain, and Yorkshire, of which Duroc and Hampshire pigs were developed mainly in North America [7]. After the long-term practice of tests for the presence of combinations of abilities, a terminal crossbreeding system with three pig breeds, namely, Duroc × (Landrace × Yorkshire) (DLY), is generally used for commercial pork production [8]. Landrace and Yorkshire pigs share prominent traits for pork production, typically, long carcass length, thin subcutaneous fat layer, large hams, good mothering ability, and high muscularity in the carcass [9,10]. Duroc pig is mainly used to enhance the growth rate and intramuscular fat in this three-way crossbreeding system [9]. The pigs generated by this system exhibit a collection of excellent traits, such as high productivity, rapid growth, and desirable pork quality and pork production [9].

Despite the above background, studies on heterosis in pigs performed to date have been limited compared with those on plants (*Arabidopsis thaliana* [11,12], maize [13], and rice [14,15]), fish [16], birds [17,18], and other mammals [19,20]. To explore the transmission of genetic information in DLY, we analyzed the whole-genome data of 30 individuals from the three pig breeds and identified 529.93 K SNPs of breed-of-origin of alleles that probably contribute to the improved crossbred performance. To explore the gene expression pattern of DLY, we carried out whole-genome sequencing of ten individuals of a DLY pig family, as well as transcriptome data of four representative tissues (including adipose, skeletal muscle, heart and liver). From these, we identified a large number of high-confidence allele-specific expression (ASE) genes, among which four ASE genes (*KMO*, *PLIN4*, *POPDC3* and *UGT1A6*) were found to be shared over all DLY pigs, suggesting that ASEs may play important roles on DLY three-way crossbreeding. Our findings are valuable for understanding the genetic architecture of DLY three-way crossbreeding, which should be beneficial for pig breeding and production.

## Results

### Simulation of three-way crossbreeding

Genomic prediction for purebred pigs regarding their crossbred performance can be based on estimation of the effects of SNPs from purebreds on the performance of their crossbred offspring [21,22]. For crossbred performance, SNP effects might be breed-specific due to differences between purebreds in allele frequency (AF) and linkage disequilibrium (LD) patterns between SNPs and quantitative trait loci (QTL) [21,22].

To estimate the breed-specific effects in terminal crossbred pigs of DLY, we calculated the probability of heterozygous SNP for each SNP locus of simulated offspring generated by three kinds of three-way crossbreeding systems **(Figure 1a)**. This was based on the ~12.56 million (M) SNPs of the publicly available genome sequencing data of 30 pig individuals from the three purebreds (including 11 Duroc, 9 Landrace, and 10 Yorkshire) with mean genome coverage of ~13.2× for each individual **(Table S1)**. Compared with the other two kinds of three-way crossbreeding system (YDL and LYD), DLY exhibited a significant distinct distribution of SNP number (**Figure 1b**, mean Pearson’s *r* = −0.99, *P* < 0.01), which was mainly due to more high-probability heterozygous SNPs in DLY (~949.05 K SNPs in DLY versus ~439.13 K SNPs in YDL and ~455.98 K SNPs in LYD when the probability of heterozygous SNP was higher than 60%; ~14.85 K SNPs in DLY versus 199 SNPs in YDL and 549 SNPs in LYD when the probability of heterozygous SNP was higher than 90%).

**Figure 1.**
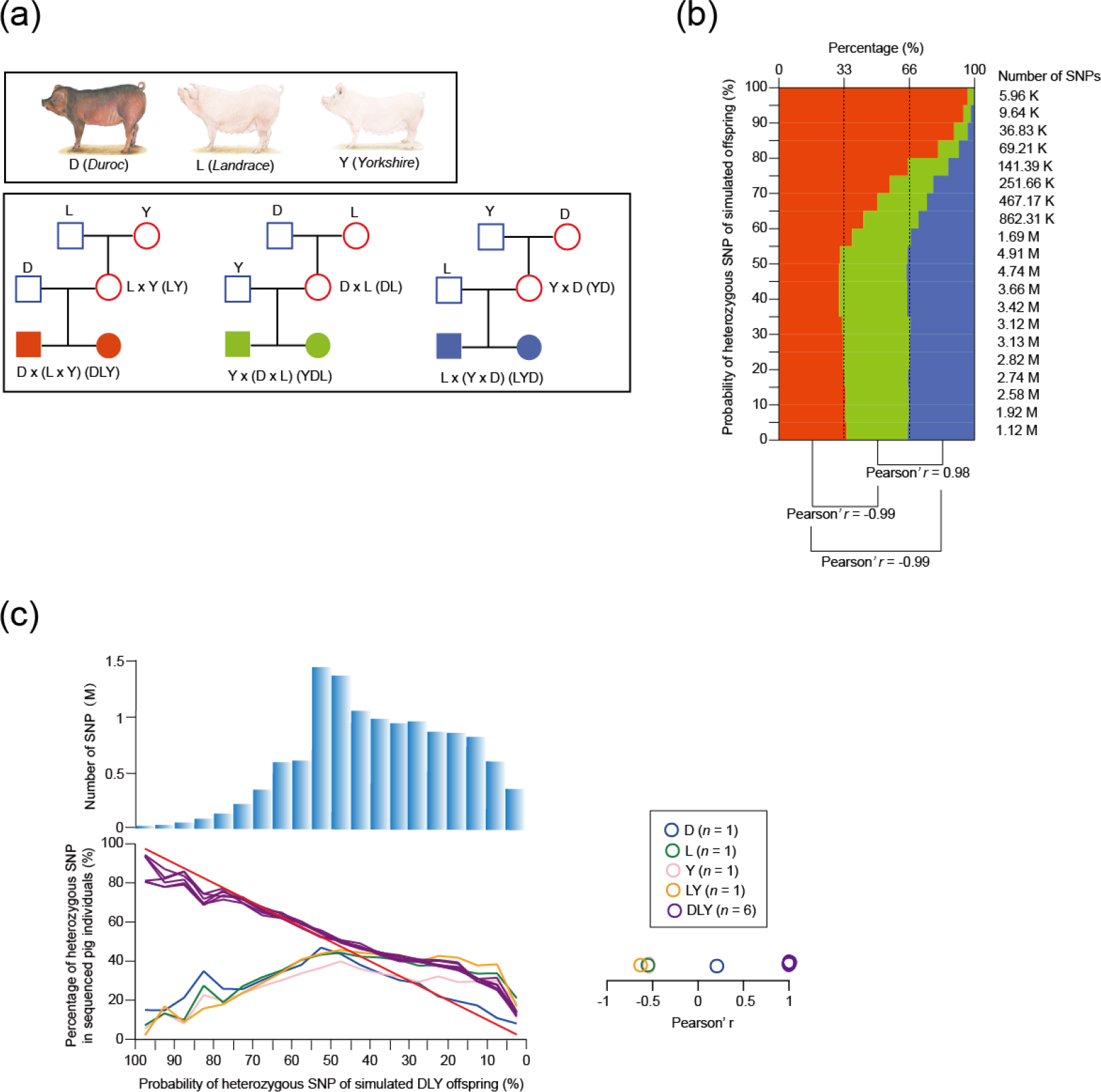
Simulation of three-way crossbreeding and validation of the accuracy of the probability of heterozygous SNP. (a) Three kinds of simulated three-way crossbreeding system among Duroc, Landrace, and Yorkshire. We did not consider the influence of sex chromosomes (X and Y) on crossbreeding, so each simulated three-way crossbreeding system represented two forms [e.g., D × (L × Y) is equivalent to D × (Y × L)]. (b) SNP number statistics of simulated offspring of three kinds of three-way crossbreeding for 20 equal intervals (based on the probability of a heterozygous SNP, from 0% to 100%, with intervals of 5%). Pearson’s r was inferred between each of two kinds of three-way crossbreeding. (c) Percentages of heterozygous SNPs of the sequenced ten pig individuals for 20 equal intervals (based on the probability of heterozygous SNP of simulated DLY offspring, from 0% to 100%, with intervals of 5%) were calculated. Pearson’s r was inferred for each comparison between simulated DLY offspring (red line) and sequenced pig individual.

To confirm the accuracy of the probability of heterozygous SNP simulated by our method, we sequenced a DLY pig family [including two grandparents (a male Landrace and a female Yorkshire), two parents (a male Duroc and a female LY crossbred pig), and six offspring (three male and three female DLY crossbred pigs)] **(Figure S2a)** and identified a total of ~14.12 M SNPs. This comprehensive collection of variations covered ~74.11% of the SNPs identified from the three downloaded pig breeds, and consistent with ~98.22% of the homozygous SNPs and ~98.36% of the heterozygous SNPs identified from Illumina’s Porcine 60K Genotyping Bead- Chip (v.2) for each individual (**Tables S2** and **S3**). The percentage of heterozygous SNPs of each pig individual was calculated for 20 equal intervals (based on the probability of heterozygous SNP of simulated DLY offspring, from 0% to 100%, at intervals of 5%) to make a comparison with the expected percentage of heterozygous SNPs of simulated DLY offspring **(Figure 1c)**. As expected, the distribution was significantly similar between the six DLY pig individuals and simulated DLY offspring (mean Pearson’s *r* = 0.98, *P* < 0.01), but significantly different between the other four pig individuals and simulated DLY offspring (mean Pearson’s *r* = −0.39, *P* < 0.01) (**Figure 1c**). This strongly confirmed the accuracy of the probability of heterozygous SNP simulated by our method.

The result presented above was most likely contributed to the smaller genetic divergence between L and Y **(Figure S1a** and **S1b)**, which suggests that they may share a more recent ancestor. As a result, the simulated offspring of LY exhibited a significant distinct distribution of SNP number (**Figure S1d**, mean Pearson’s *r* = −0.93, *P* < 0.01) compared with the other two kinds of two-way crossbreeding (DL and DY, **Figure S1c**), which was mainly due to fewer high-probability heterozygous SNPs in LY (~641.27 K SNPs in LY versus ~1.37 M SNPs in DL and ~1.27 M SNPs in YD when the probability of heterozygous SNPs was higher than 60%; 1,211 SNPs in LY versus ~43.62 K SNPs in DL and ~36.83 K SNPs in YD when the probability of heterozygous SNPs was higher than 90%). Taking these findings together, we suggest that DLY is a more effective strategy for three-way crossbreeding among Duroc, Landrace, and Yorkshire breeds as it could generate a large number of stably inherited heterozygous SNPs.

### Identification of breed-of-origin of alleles

To identify the breed-of-origin of alleles that most likely contribute to the prominent performance of crossbred individuals, we separately quantified the genetic differentiation (*F*_st_) for each SNP locus between D and L (D-L) as well as D and Y (D-Y). We used an empirical procedure and selected SNP loci simultaneously exhibiting a significant high value of *F*_st_ (the top 5% right tail, where *F*_st_ was larger than 0.61) and a significant high value of probability of heterozygous SNP of simulated DLY offspring (the top 5% right tail, where the probability of heterozygous SNP was higher than 0.64) of the empirical distribution (**Figure 2a** and **2b**). Using this approach, we identified ~399.48 K and ~379.21 K SNPs of candidate breed-of-origin of alleles for D-L and D-Y, respectively (**Figure 2a** and **2b**), which constituted a final combined collection of 529.93 K SNPs of breed-of-origin of alleles **(Figure 2c)**. As expected, these identified SNPs of breed-of-origin of alleles exhibited a significant higher percentage of heterozygous SNPs (~69.52%) than the genomic background (~47.54%) in the six DLY pig individuals (**Figure 2d**, *P* < 1.38 × 10^−6^, paired *t*-test).

**Figure 2.**
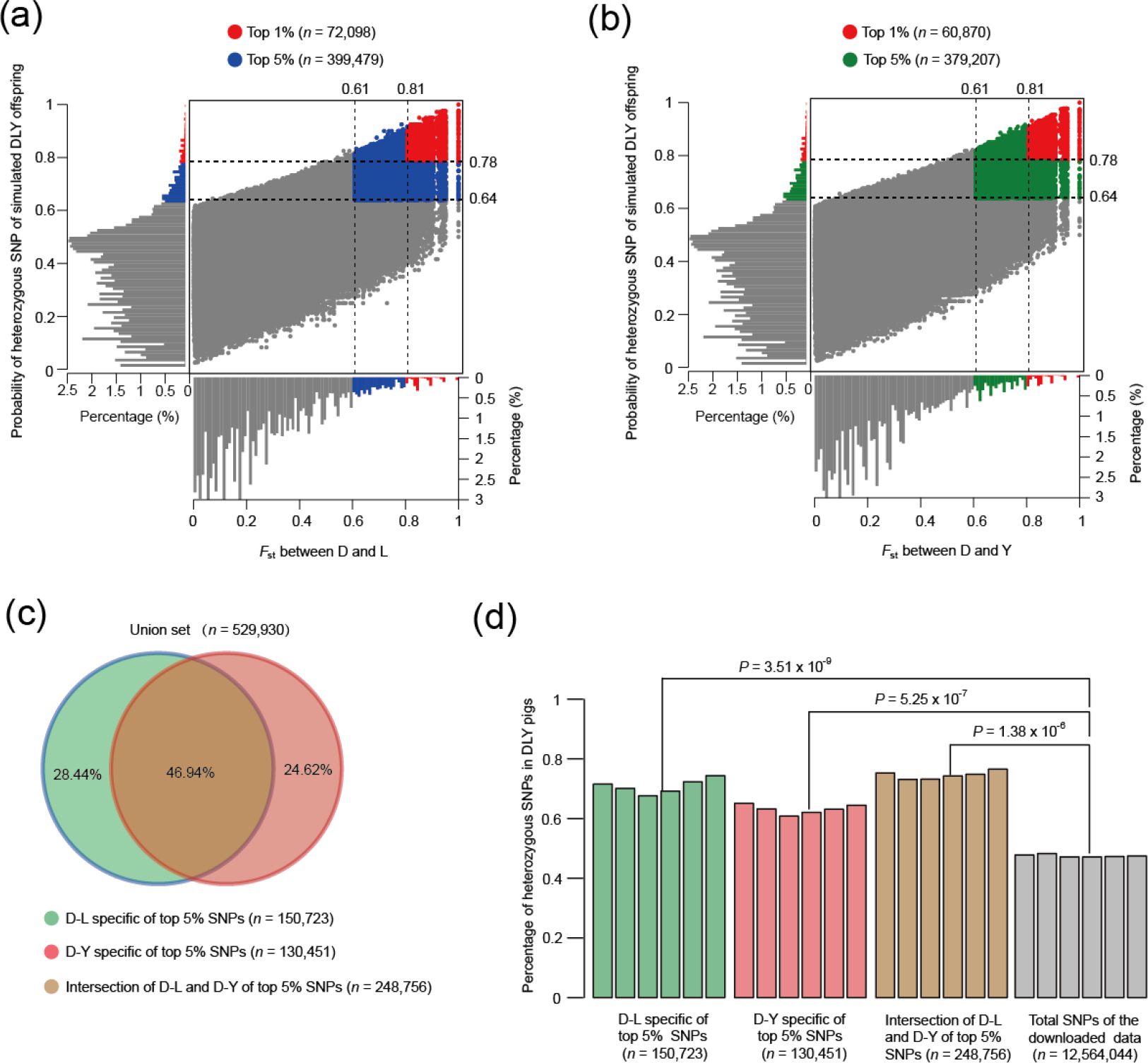
Identification of breed-of-origin of alleles. (a) Distribution of probability of heterozygous SNP of simulated DLY offspring and *F*_st_ value between D and L for each SNP locus. A total of ~399.48 K SNPs of candidate breed-of-origin of alleles were selected that simultaneously showed a significant high probability of heterozygous SNP (the top 5% right tail, where the probability of heterozygous SNP was larger than 0.64) and a significant high *F*_st_ value (the top 5% right tail, where *F*_st_ value was larger than 0.61). (b) Distribution of probability of heterozygous SNP of simulated DLY offspring and *F*_st_ value between D and Y for each SNP locus. A total of ~379.21 K SNPs of candidate breed-of-origin of alleles were selected that simultaneously showed a significant high probability of heterozygous SNP (the top 5% right tail, where the probability of heterozygous SNP was larger than 0.64) and a significant high *F*_st_ value (the top 5% right tail, where *F*_st_ value was larger than 0.61). (c) The full collection of 529.93 K SNPs of breed-of-origin of alleles combining candidate breed-of-origin of alleles of D-L and candidate breed-of-origin of alleles of D-Y. (d) Paired *t*-test was used to compare the percentage of heterozygous SNPs of breed-of-origin of alleles and the genomic background in the six DLY pig individuals.

Although we observed similar occurrences between breed-of-origin of alleles and the genomic background (**Table S4**, Pearson’s *r* = 0.99, *P* < 0.01) for five different genomic elements (i.e., upstream, downstream, intergenic, intronic, and exonic), we also identified a large number of functional categories (GO and KEGG) for SNPs of breed-of-origin of alleles located in upstream (**Table S5**), downstream (**Table S6**), exonic (**Table S7**), and intronic regions (**Table S8**), some of which were strongly associated with growth rate (epidermal growth factor receptor signaling pathway, *P* = 0.034; cartilage development, *P* = 4.1 × 10^−3^; skeletal system morphogenesis, *P* = 4.9 × 10^−3^; regulation of cell proliferation, *P* = 0.029; post-embryonic development, *P* = 0.048), disease resistance (MHC class II receptor activity, *P* = 1.1 × 10^−3^; response to antibiotics, *P* = 0.029; antigen processing and presentation of exogenous peptide antigen via MHC class II, *P* = 3.8 × 10^−3^; inflammatory mediator regulation of TRP channels, *P* = 4.9 × 10^−4^; T-cell costimulation, *P* = 0.048), pork production (lipid catabolic process, *P* = 6.9 × 10^−3^; acyl-CoA hydrolase activity, *P* = 0.047; regulation of fat cell differentiation, *P* = 0.041; long-chain fatty-acyl-CoA biosynthetic process, *P* = 0.016; regulation of lipolysis in adipocytes, *P* = 0.025), and reproductive performance (oocyte meiosis, *P* = 0.026; ovarian steroidogenesis, *P* = 0.013; oxytocin signaling pathway, *P* = 5.1 × 10^−4^).

To test the influence of the breed-of-origin of alleles on protein-coding genes, we selected 50 nonsynonymous SNPs of breed-of-origin of alleles simultaneously exhibiting the highest values of probability of heterozygous SNP and high-confidence heterozygous SNPs in DLY individuals (five or more heterozygous SNPs for six DLY pig individuals) **(Figure 3a)**. These selected nonsynonymous SNPs correspond to 50 genes that were supposed to have close associations with the improved performance of crossbred individuals **(Figure 3a)**. A representative example of these is *AKAP9* (A-kinase anchoring protein 9), which is a member of structurally diverse proteins that have the common function of binding to the regulatory subunit of protein kinase A (PKA) [23, 24]. The detail information of variant in *AKAP9* was showed in **Figure 3b**. Mice with *AKAP9* knockout displayed decreased body fat and body weight, hematopoietic abnormalities, and an atypical plasma chemistry profile, suggesting that this gene may be important for fat and body weight development [25,26]. Detailed information for the other genes is presented in **Table S9**. Taken together, our findings provide a valuable resource to further explore the mechanism behind heterosis in DLY three-way crossbreeding.

**Figure 3.**
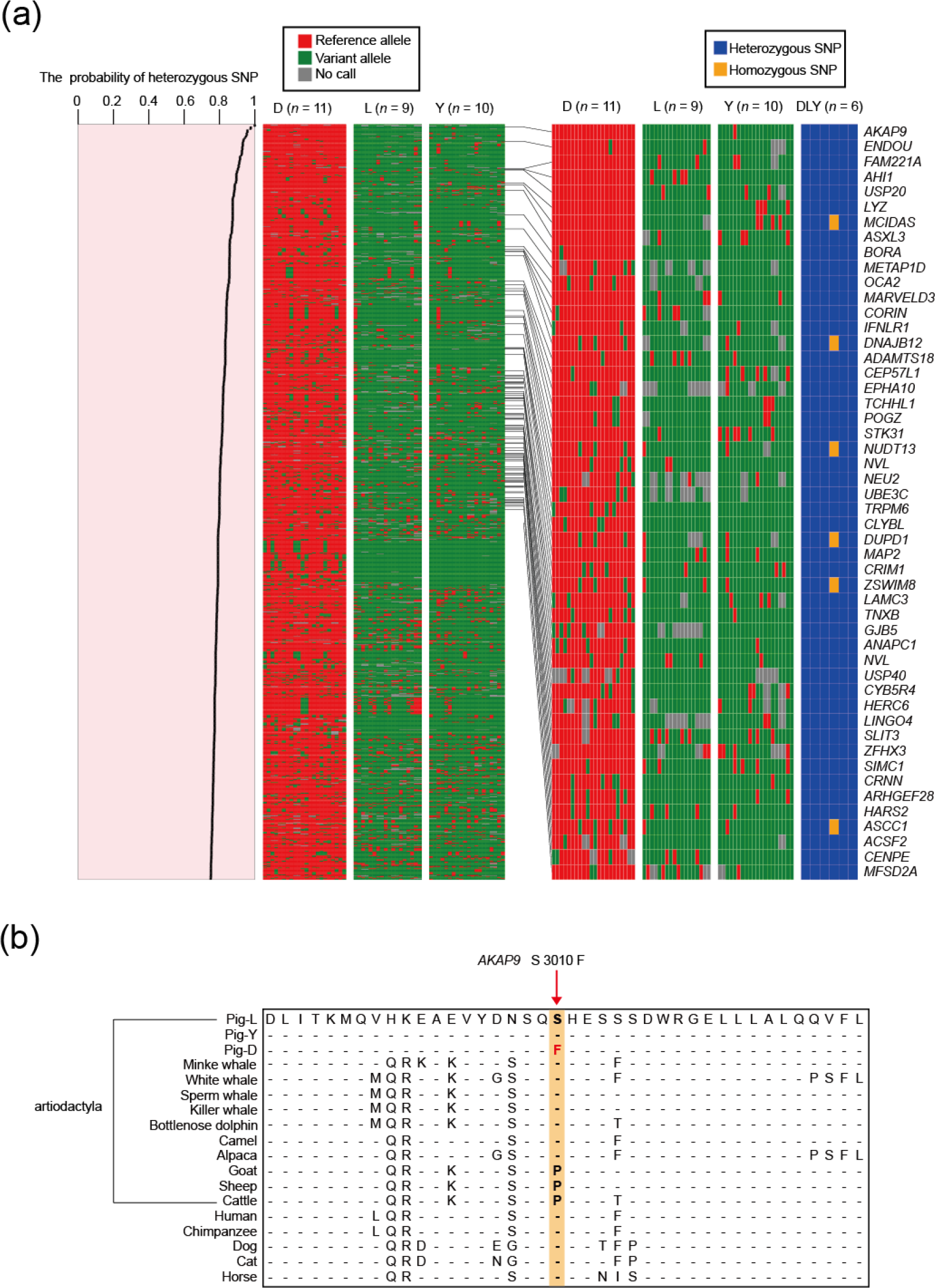
Allele distribution and genes related to nonsynonymous SNPs of breed-of-origin of alleles. (a) We showed the allele distribution among D, L, and Y for 1000 exonic SNPs of breed-of-origin of alleles with the highest probability of heterozygous SNP and selected 50 nonsynonymous SNPs of breed-of-origin of alleles that simultaneously showed the highest probability of heterozygous SNP and high-confidence heterozygous SNPs in DLY pig individuals (five or more heterozygous SNPs for six DLY pig individuals). These 50 nonsynonymous SNPs of breed-of-origin alleles correspond to 50 genes. (b) Multiple protein sequence alignment of *AKAP9* among pig and 15 other mammals. The nucleotide on the reference pig genome chr9 : 72,027,468 was identified as a breed-of-origin allele, caused an amino acid conversion from S to F in the Duroc breed.

### Allele-specific expression (ASE)

To explore the gene expression pattern during the progress of DLY three-way crossbreeding, we sequenced the transcriptome of four representative tissues, including adipose (energy metabolism), skeletal muscle (body movement and supporter), heart (hematopoiesis and circulatory), and liver (digestion and detoxification) for six DLY individuals (**Table S10**). We detected 2,071–2,261 (65.35%–71.35%) heterozygous SNPs in these six individuals overlapping with the 3,169 exonic SNPs of breed-of-origin of alleles; the majority of the overlapping heterozygous SNPs (~96.03%) could be distinguished between the paternal and maternal alleles (**Figure S3a**).

We used a strict criteria (*P*-value < 0.05 and fold change > 2) to identify high-confidence ASEs in adipose **(Figure S4)**, skeletal muscle **(Figure S5)**, heart **(Figure S6)**, and liver **(Figure S7)** and found that ASEs were significantly enriched in breed-of-origin of alleles **(Figure 4e**, mean ASE percentage of ~3.79% for breed-of-origin of alleles versus ~2.04% for the genomic background, *P* < 4.82 × 10^−3^, paired *t*-test). We compared the distribution of ASEs in six DLY pigs and found that four ASEs were shared among all DLY pigs, while the others were individual-specific **(Figure 5a)**. We then focused on these four ASEs to explore the influence of ASEs on DLY three-way crossbreeding system.

**Figure 4.**
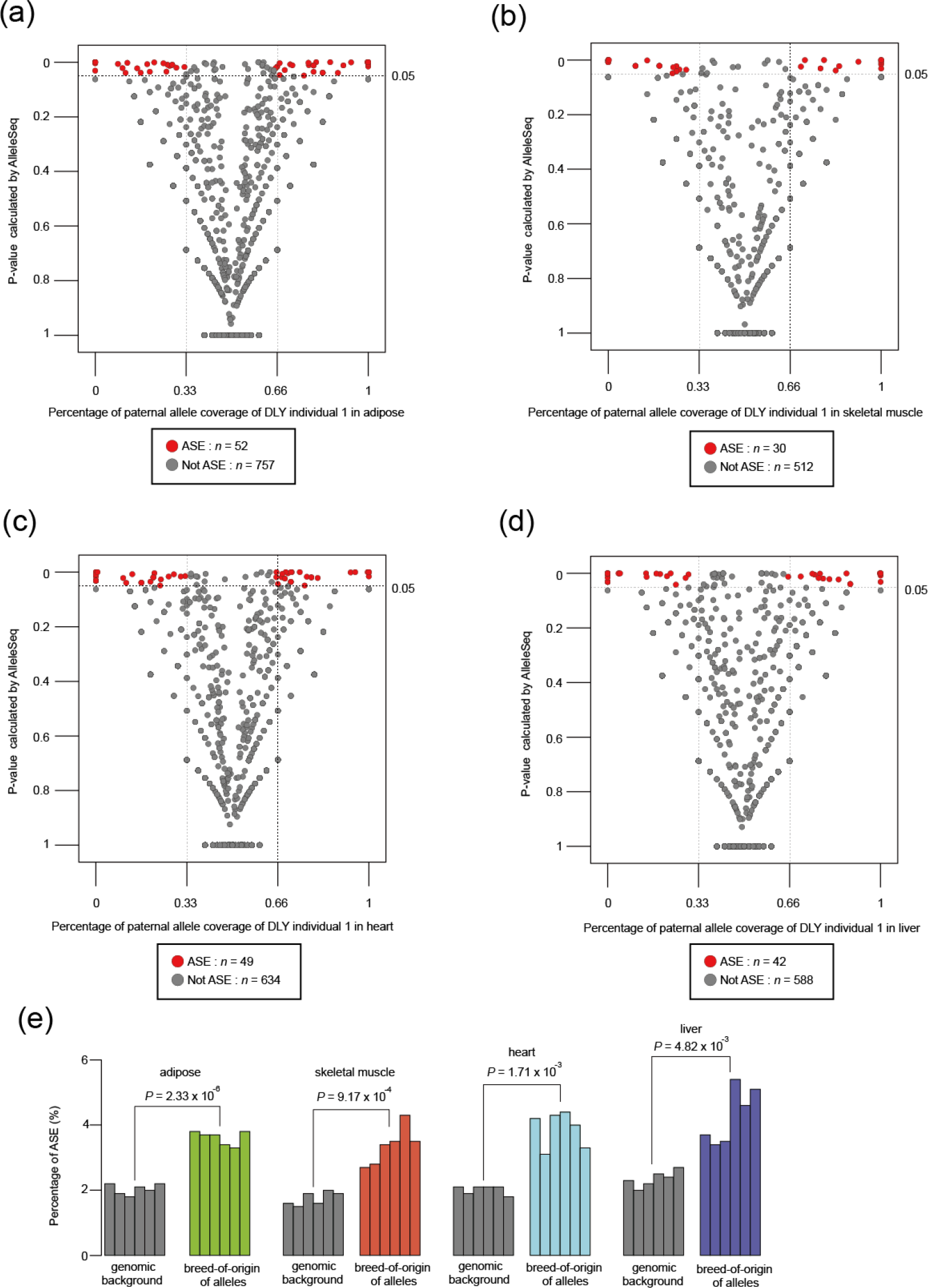
Identification of allele-specific expression (ASE). (a) ASE identification of adipose for DLY individual 1. (b) ASE identification of skeletal muscle for DLY individual 1. (c) ASE identification of heart for DLY individual 1. (d) ASE identification of liver for DLY individual 1. (e) Paired *t*-test was used for comparison of ASE occurrence between breed-of-origin of alleles and the genomic background for adipose, skeletal muscle, heart, and liver.

**Figure 5.**
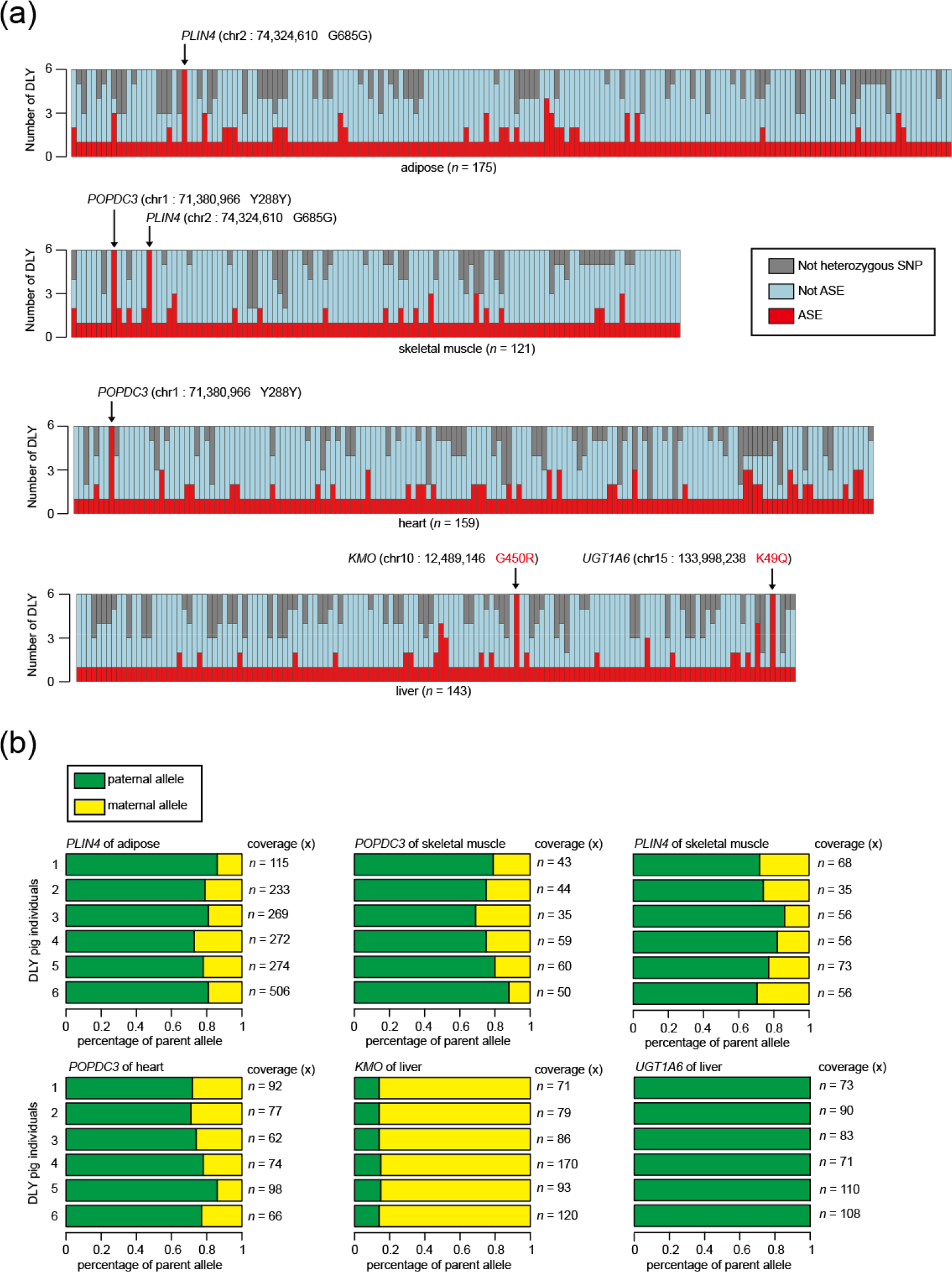
ASE distribution over six DLY pig individuals. (a) We analyzed the ASE distribution over the six DLY pig individuals for adipose, skeletal muscle, heart, and liver. All SNP loci were classified into three categories [not heterozygous SNP (gray), not ASE (light blue), and ASE (red)]. (b) Distribution of the percentages of paternal allele (green) and maternal allele (yellow) for the four shared ASEs over six DLY pig individuals.

*KMO* (kynurenine 3-monooxygenase) contained a nonsynonymous SNP (position 12,489,146 on chromosome 10), which was detected to express significant more maternal allele (~85.46%) than paternal allele (~14.54%) in liver of all DLY individuals (**Figure 5b**, *P* < 10^−16^, paired *t*-test). *KMO* encodes a mitochondrial outer-membrane protein that catalyzes the hydroxylation of an L-tryptophan metabolite, L-kynurenine, to form L-3-hydroxykynurenine; recent studies have also suggested that kynurenine-3-monooxygenase deficiency is associated with disorders of the brain (e.g., schizophrenia and tic disorders) and of the liver [27,28,29]. *PLIN4* (perilipin 4) contained a synonymous SNP (position 74,324,610 on chromosome 2), which was detected to express significant more paternal allele (~79.27%) than maternal allele (~20.73%) in adipose (*P* < 3.5 × 10^−10^, paired *t*-test) as well as significant more paternal allele (~75.87%) than maternal allele (~24.13%) in skeletal muscle (*P* < 2.3 × 10^−5^, paired *t*-test) of all DLY individuals (**Figure 5b**). *PLIN4* is one of the perilipin members of the PATS family, which share high amino acid sequence similarity and associate with lipid droplets [30]. It may have a close association with insulin resistance and obesity risk due to its protection of lipid droplets from lipases [31]. Studies have revealed close relationships between perilipin genes and body weight and body composition during weight maintenance [32], suggesting that *PLIN4* plays an important role in adipose storage and development for DLY crossbred pigs. *POPDC3* (popeye domain containing 3) contained a synonymous SNP (position 71,380,966 on chromosome 1), which was detected to express significant more paternal allele (~78.01%) than maternal allele (~21.99%) in skeletal muscle (*P* < 4.5 × 10^−8^, paired *t*-test) as well as significant more paternal allele (~76.55%) than maternal allele (~23.45%) in heart (*P* < 1.7 × 10^−8^, paired *t*-test) of all DLY individuals (**Figure 5b**). *POPDC3* encodes a member of the POP family of proteins containing three putative transmembrane domains [33]. Although the function of this gene is not very clear, it is predominantly expressed in skeletal and cardiac muscle, suggesting that this gene has an important function in these tissues [34]. *UGT1A6* (UDP glucuronosyltransferase family 1 member A6) contained a nonsynonymous SNP (position 133,998,238 on chromosome 15), which was detected to specifically express paternal allele in liver of all DLY individuals (**Figure 5b**). *UGT1A6* is related to the function of the liver due to its function of transforming small lipophilic molecules (steroids, bilirubin, hormones, and drugs) into water-soluble, excretable metabolites [35,36], suggesting this an important role associated with liver. Taking these findings together, we suggest that ASEs may also play important roles in improving the performance of DLY crossbred pigs.

## Discussion

In this study, we investigated the DLY three-way crossbreeding system via genomic analyses. Based on ~12.56 M population-scale SNPs (30 individuals from three pig breeds), ~14.12 M family-scale SNPs (including ten pig individuals), and transcriptome data (including adipose, skeletal muscle, heart, and liver), we identified 529,930 breed-of-origin of alleles and numerous high-confidence ASEs, including four shared ASE genes (*KMO*, *PLIN4*, *POPDC3*, and *UGT1A6*). These findings should be valuable for pig breeding and production in the future and provide an excellent model of crossbreeding analysis for many other agricultural animals. We expect that the application of more resources and technologies will reveal more comprehensive and significant discoveries about genetic mechanism of heterosis for pigs in the future.

## Materials and methods

### Genome sequencing data of pigs

In this study, we downloaded genome sequencing data of 30 pig individuals (including 11 Duroc pigs, 9 Landrace pigs and 10 Yorkshire pigs) from NCBI (Supplementary URLs) **(Table S1)**. To produce a tidy dataset for population variants calling, we randomly selected 33 Gb (~13.2× coverage) data for each pig individual **(Table S1)**. We also sequenced a DLY pig family including 10 pig individuals **(Figure S2a)** using Illumina Hiseq 4000 platform. In total, we generated ~962 Gb paired-end 100-bp (PE100) sequencing data for 10 pig individuals, with an average genome coverage of ~38.48× **(Table S2)**.

### Mapping and SNP calling

The genome sequencing data of pigs in this study were first mapped to reference pig genome (version Sus_scrofa.11.1, Supplementary URLs) using BWA (version 0.7.8) [37] with default options. We used “MarkDuplicates” module in package Picard (version 1.48, Supplemental URLs) to remove duplicated reads. The module “HaplotypeCaller” in Genome Analysis Toolkit (GATK; version 3.7) [38] was used to identity SNPs, which were next filtered by following criteria: QUAL < 30.0, QD < 2.0, MQ < 40.0, FS > 60.0. Additionally, we required genome coverage equal or larger than 4 and SNP integrity equal or larger than 0.7, which were analyzed by VCFtools (version 1.15) [39]. Finally, we detected a total of ~12.56 M SNPs of the three downloaded pig breeds **(Table S1)** and ~14.12 M SNPs of the DLY pig family **(Table S2)**. We used Illumina’s porcine 60K Genotyping Bead-Chip (v.2) to validate the accuracy of SNP identification for each individual of DLY pig family **(Table S3)**. To test whether the 10 individuals of DLY pig family had a correct sibship, we used PLINK (version 1.07) [40] to calculate PI_HAT value between two different individuals using the ~14.12 M family-scale SNPs **(Figure S2b)**. The expected value of PI_HAT is 0.5 between offspring and parents and decreased to 0.25 between grandparents and offspring. If two pig individuals shared the same parents, the value of PI_HAT is close to 0.5. The extreme value of 0 means two individuals have no sibship. Based on this method, we verified the correct sibship of the sequenced DLY pig family **(Figure S2b)**.

### Calculation of the probability of heterozygous SNP

To calculate the probability of heterozygous SNP of simulated offspring generated by different crossbreeding systems (including two-way and three-way crossbreeding), we calculated frequency of three genotypes (homozygous identical with reference, homozygous distinct from reference, heterozygous) of the ~12.56 M population-scale SNPs for Duroc (11 individuals), Landrace (9 individuals) and Yorkshire (10 individuals) respectively. Based on the above results, we calculated the frequency of the same three genotypes for the simulated offspring generated by three kinds of two-way crossbreeding systems (DL, LY and YD, **Figure S2c**). Similarly, the frequency of the same three genotypes of simulated offspring generated by three kinds of three-way crossbreeding systems **(Figure 1a)** were inferred using same method.

### Population genetic analyses

The phylogenetic tree was inferred using the package treebest (version 1.9.2, Supplementary URLs) under the p-distances model using the ~12.56 M population-scale SNPs with a bootstrap of 100. We performed PCA (principal component analysis) with software GCTA (version 1.91.3beta, Supplementary URLs) using the ~12.56 M population-scale SNPs. *F*_st_ value for each SNP locus was calculated by VCFtools (version 1.15) [39] between D and L as well as D and Y respectively.

### Annotation of breed-of-origin of alleles

We performed annotation for SNPs of breed-of-origin of alleles with ANNOVAR (version 3.5c) [41]. The current pig gene annotation file (version Sus_scrofa.11.1.93, Supplementary URLs) was used to build ANNOVAR database of pig and SNPs of breed-of-origin of alleles were classified into 5 different genomic elements, including upstream, downstream, exonic, intronic and intergenic regions **(Table S4)**.

### Function enrichment analyses

We performed function enrichment analyses with DAVID (Database for Annotation, Visualization and Integrated Discovery, version 6.8) [42]. We prefer to use human annotation database rather than pig so we downloaded ortholog gene information between pig and human from Ensembl genome browser website (Supplementary URLs) and translated pig genes to human ortholog genes.

### Allele-specific expression (ASE) analysis

To identify allele-specific expression (ASE) during the progress of DLY three-way crossbreeding system, we sequenced 4 representative tissues, including adipose (subcutaneous adipose), skeletal muscle (longissimus dorsi muscle), heart and liver, generating a total of ~111.5 Gb transcriptome data **(Table S8)**. We used software vcf2diploid (version 0.2.6a, Supplementary URLs) to create paternal genome and maternal genome based on the ~14.12 M family-scale SNPs for the 6 DLY pig individuals respectively. Transcriptome data were mapped to paternal and maternal genome with bowtie (version 1.1.2) [43] and AlleleSeq software (version 1.2a) [44] was used to calculate the *P*-value for imbalanced expression level. We identified a high-confidence ASE simultaneously with significant *P*-value smaller than 0.05 and significant percentage of paternal allele coverage smaller than 0.33 or larger than 0.66 (also means fold-change > 2).

## Data access

The whole-genome sequencing data of ten pig individuals and transcriptome data the six DLY pig individuals of the DLY family are accessible at NCBI BioProject under accession number of PRJNA507853. The genotyping data of the Illumina’s porcine 60K Genotyping Bead- Chip (v.2) have been submitted to the NCBI Gene Expression Omnibus (GEO) under accession number of GSE123327.

## FUNDING

This work was supported by grants from the National Key R & D Program of China (2018YFD0500403), the National Natural Science Foundation of China (31772576, 31522055 and 31802044), the Sichuan Province & Chinese Academy of Science of Science & Technology Cooperation Project (2017JZ0025), the Science & Technology Support Program of Sichuan (2016NYZ0042 and 2017NZDZX0002), the Earmarked Fund for China Agriculture Research System (CARS-35-01A).

### URLs

NCBI,https://www.ncbi.nlm.nih.gov/; Reference pig genome, ftp://ftp.ensembl.org/pub/release92/fasta/susscrofa/dna/Sus_scrofa.Sscrofa11.1.dna.toplevel.fa.gz/; Picard, http://picard.sourceforge.net/; treebest, http://treesoft.sourceforge.net/treebest.shtml/; GCTA, http://www.gcta-ga.org/; Reference pig annotation file, ftp://ftp.ensembl.org/pub/release93/gtf/sus_scrofa/Sus_scrofa.Sscrofa11.1.93.gtf.gz; Ensembl genome browser, http://asia.ensembl.org/index.html; vcf2diploid, http://alleleseq.gersteinlab.org/tools.html.

## Supporting information

