## Supplementary material for "Genomic analyses provide insights into genetic architecture of three-way crossbred pigs"

**Figures**

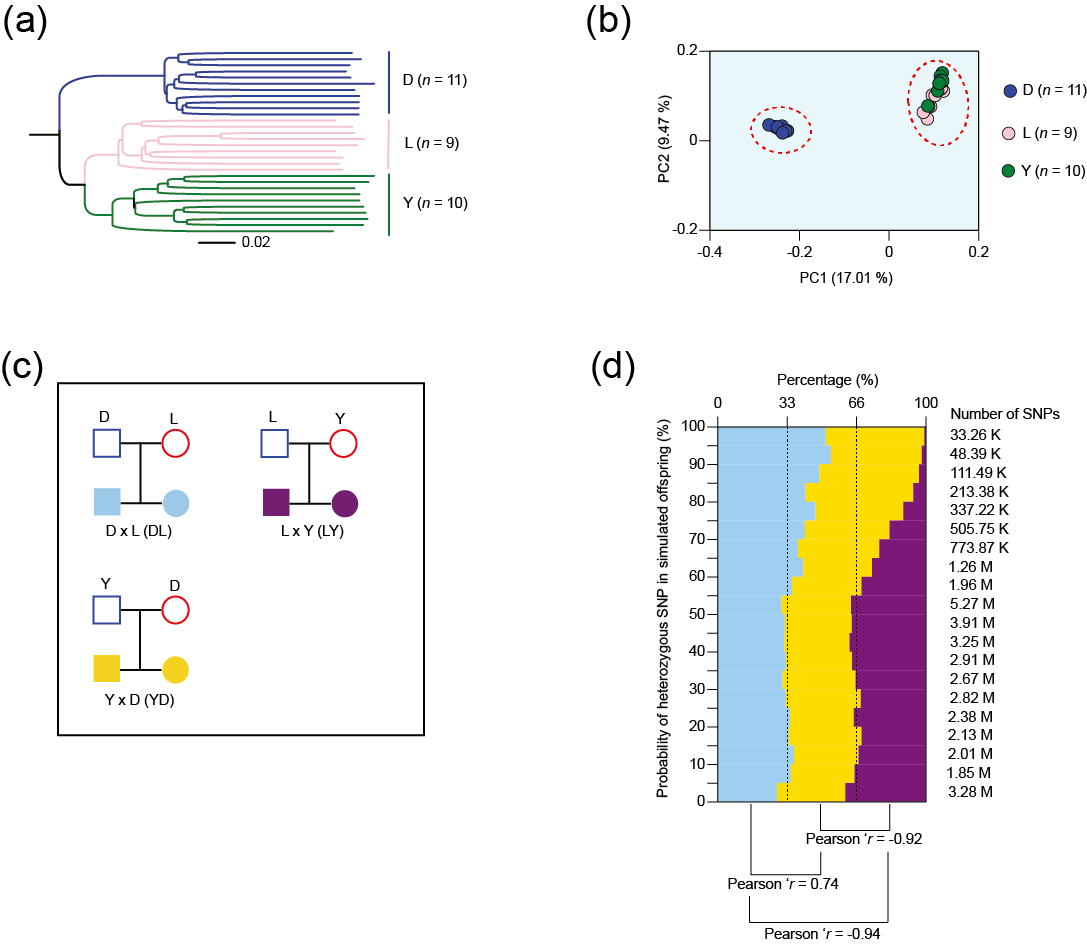

**Figure S1. Population genetic analyses and simulation of heterozygous SNP distribution for two-way crossbreeding.**

(a) Phylogenetic tree of the downloaded three pig breeds using identified ~12.56 M SNPs.

(b) Principal component analysis (PCA) of the downloaded three pig breeds using identified ~12.56 M SNPs.

(c) Simulation of heterozygous SNP distribution for three kinds of two-way crossbreeding. Similarly, we didn’t consider sex chromosome (X chromosome and Y chromosome) for analysis so each two-way crossbreeding represented two forms [e.g. D x L (DL) is equivalent to L x D (LD)].

(d) SNP number statistics of the simulated offspring of three kinds of two-way crossbreeding for 20 equal intervals (based on the probability of heterozygous SNP, from 0% to 100%, with intervals of 5%). Pearson’s r was inferred between each of two kinds of two-way crossbreeding

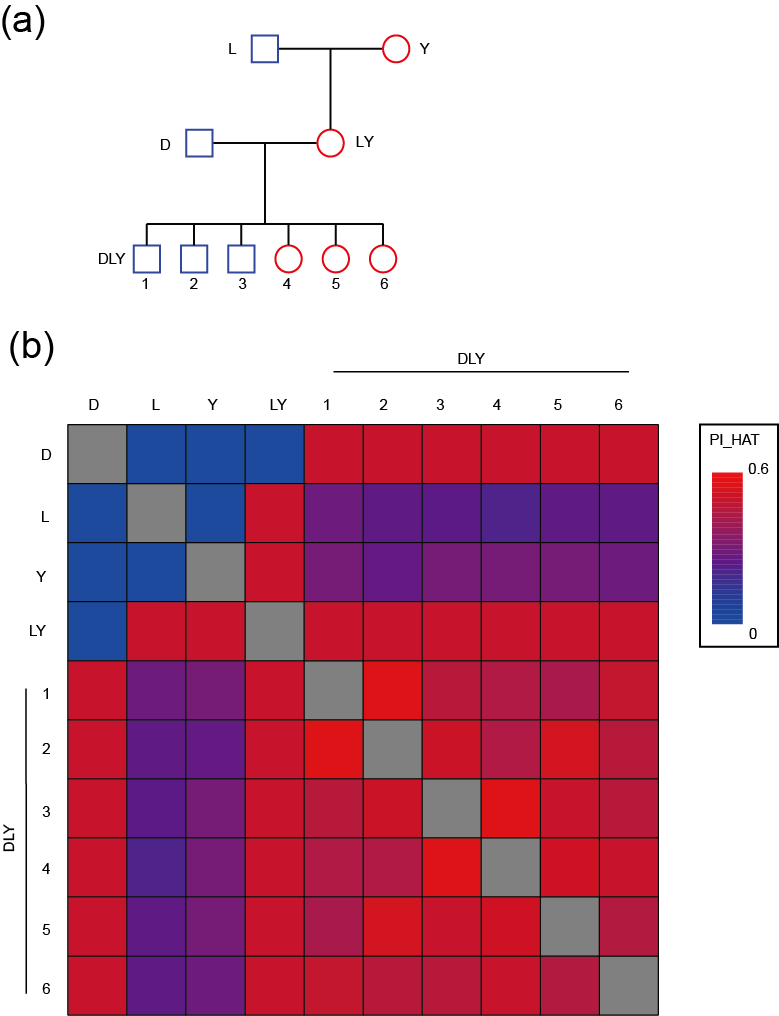

**Figure S2. Members of the DLY pig family and paternity test.**

(a) The ten pig individuals of the sequenced DLY pig family.

(b) We used PLINK to calculate PI_HAT value between each of two pig individuals based on ~14.12 M family-scale SNPs. The expected value of PI_HAT is 0.5 between offspring and parents and decreased to 0.25 between grandparents and offspring. If two pig individuals shared the same parents, the value of PI_HAT is close to 0.5. The extreme value of 0 means two individuals have no sibship.

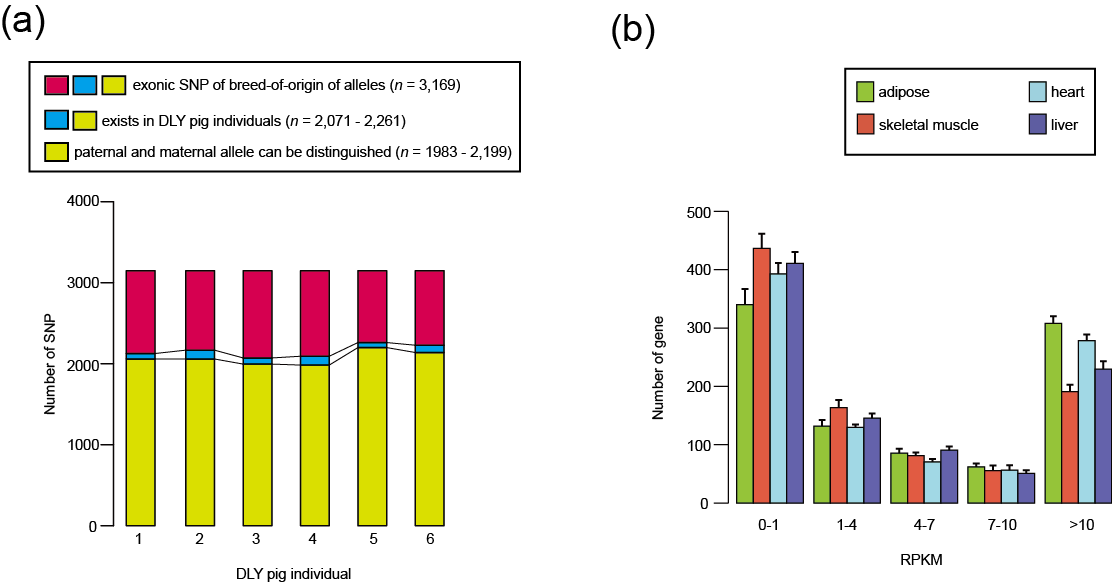

**Figure S3. Heterozygous SNP distribution and expression pattern.**

(a) Summary of the overlap between exonic SNPs of breed-or-origin of alleles and heterozygous SNPs in DLY individuals. We observed a high percentage for exonic SNPs of breed-of-origin of alleles exists in DLY pigs (65.35%-71.35%) and a high percentage of distinguishable heterozygous SNPs of DLY individuals of paternal allele and maternal allele (~96.03%).

(b) We displayed the gene expression (RPKM) of genes containing distinguishable heterozygous SNPs in adipose, skeletal muscle, heart and liver. A quite percentage of genes (34.84-0.53%) exhibited low expression pattern (RPKM < 1), which were ignored for further analysis.

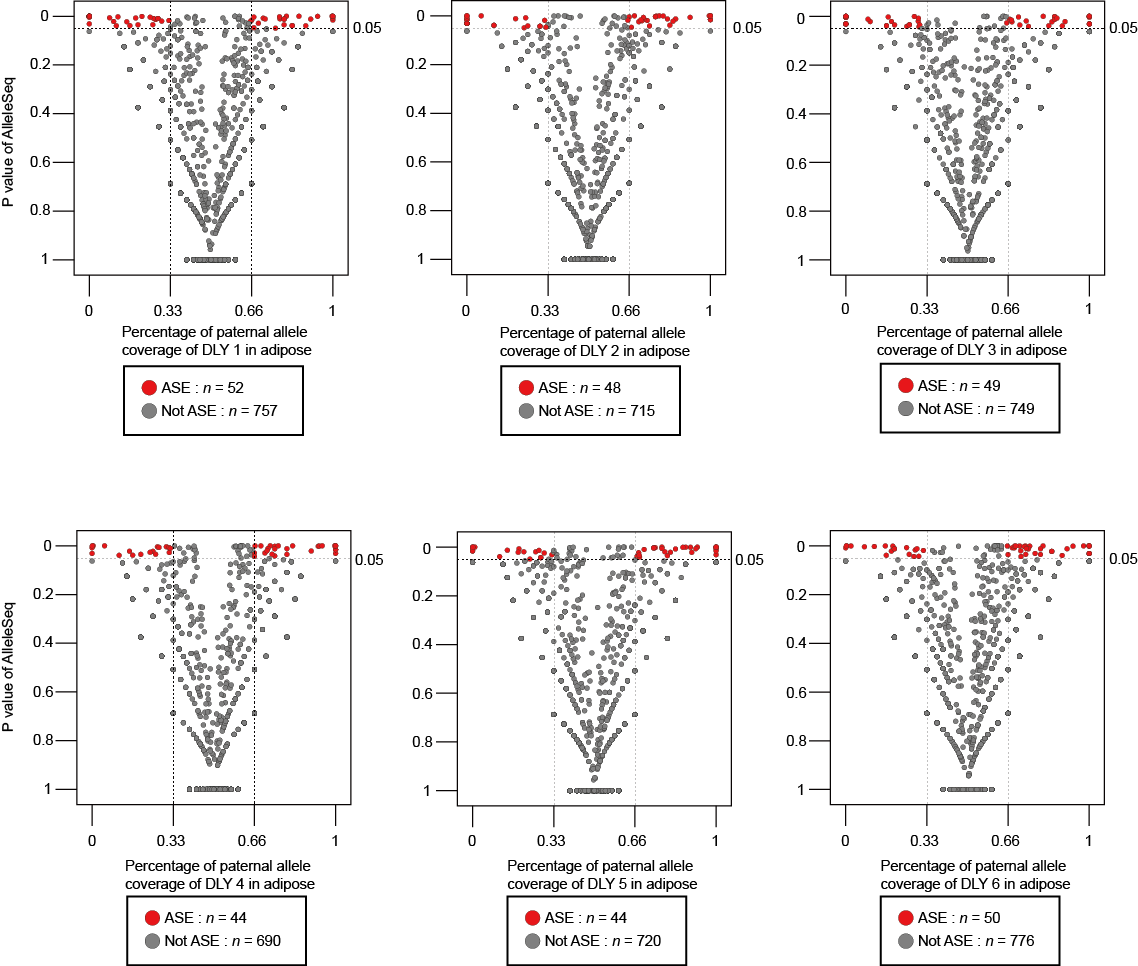

**Figure S4. ASE identification of adipose for 6 DLY individuals.**

we used a strict criteria to identify high-confidence ASEs simultaneously with significant *P*-value smaller than 0.05 and significant percentage of paternal allele coverage smaller than 0.33 or larger than 0.66 (also means fold-change > 2).

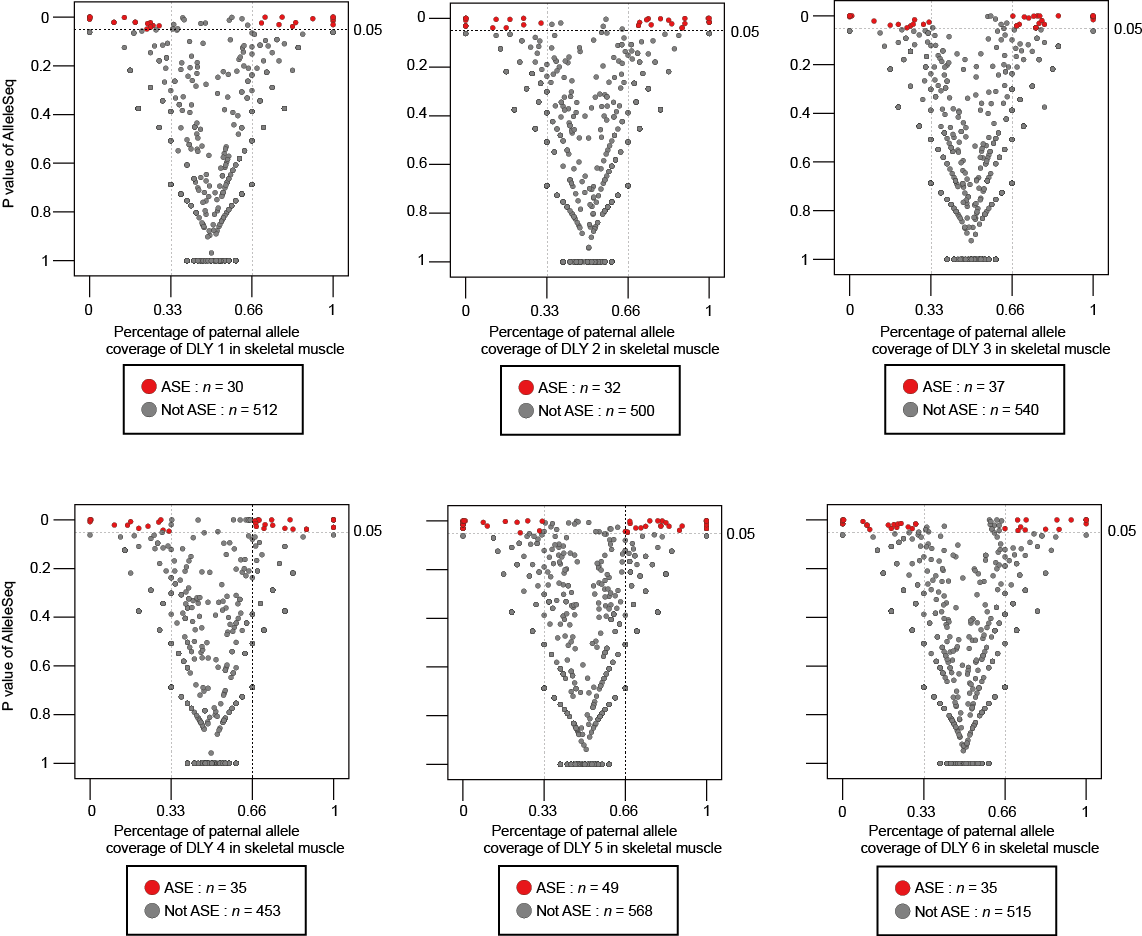

**Figure S5. ASE identification of skeletal muscle for 6 DLY individuals.**

we used a strict criteria to identify high-confidence ASEs simultaneously with significant *P*-value smaller than 0.05 and significant percentage of paternal allele coverage smaller than 0.33 or larger than 0.66 (also means fold-change > 2).

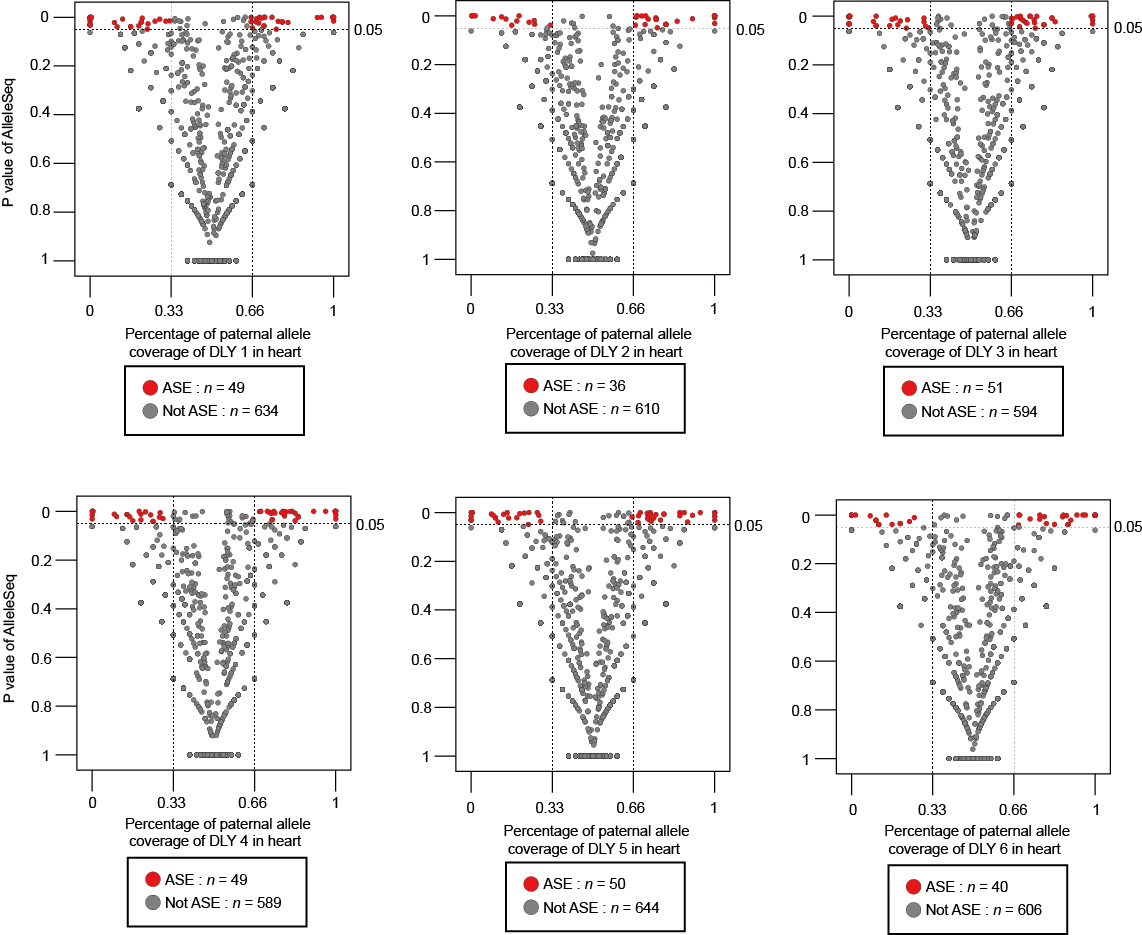

**Figure S6. ASE identification of heart for 6 DLY individuals.**

we used a strict criteria to identify high-confidence ASEs simultaneously with significant *P*-value smaller than 0.05 and significant percentage of paternal allele coverage smaller than 0.33 or larger than 0.66 (also means fold-change > 2).

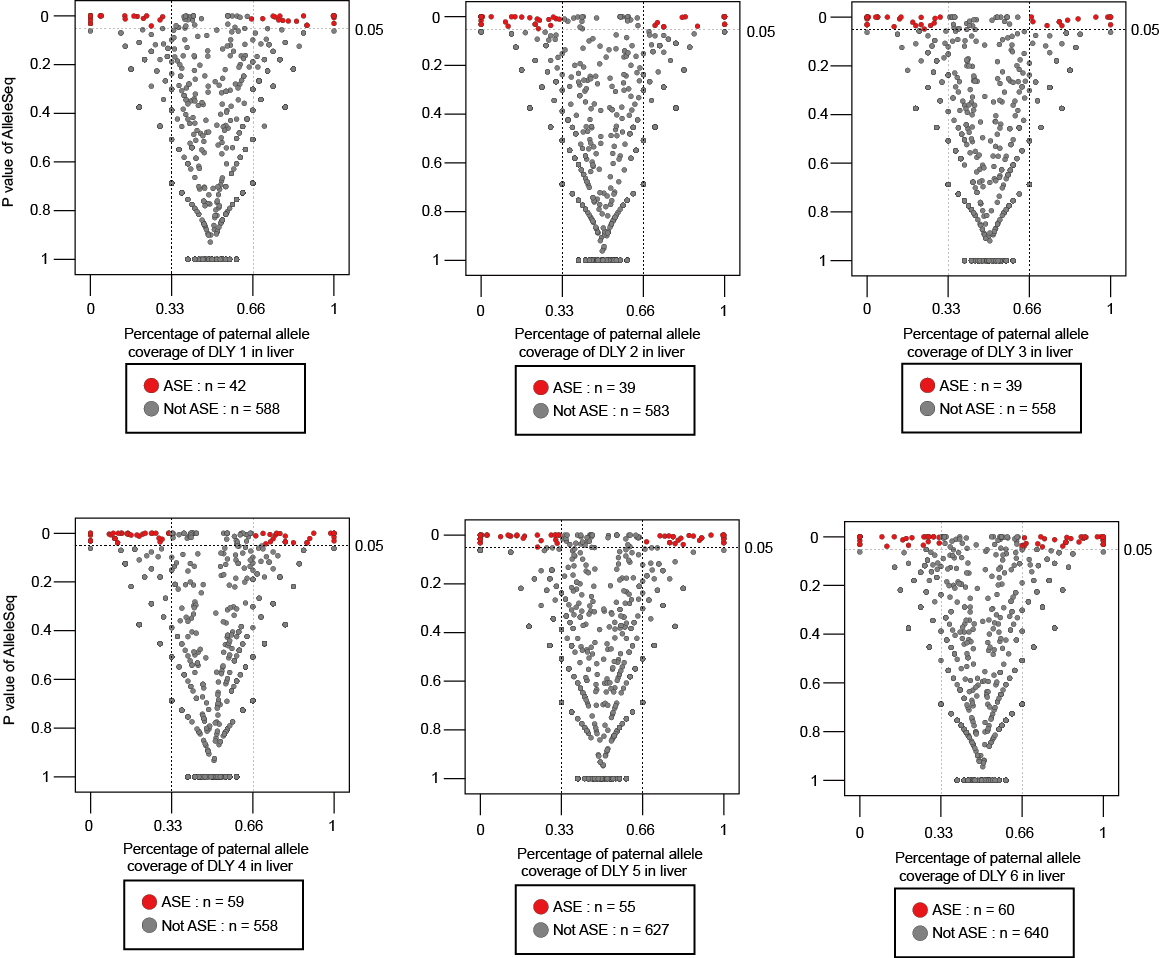

**Figure S7. ASE identification of liver for 6 DLY individuals.**

we used a strict criteria to identify high-confidence ASEs simultaneously with significant *P*-value smaller than 0.05 and significant percentage of paternal allele coverage smaller than 0.33 or larger than 0.66 (also means fold-change > 2).

**Tables**

Table S1 Data summary and SNP statistics of the downloaded pig breeds

| **Breed** | **Individual** | **PE length**  **(bp)** | **Accession**  **No.** | **High-quality**  **data (Gb)** | **Coverage (**×**)** | **Heterozygous**  **SNP** | **Homozygous**  **SNP** |
| --- | --- | --- | --- | --- | --- | --- | --- |
| D | 1 | 100 | SRX703502 | 33 | 13.2 | 2,970,948 | 1,970,902 |
|  | 2 | 100 | SRX703503 | 33 | 13.2 | 2,915,641 | 1,961,520 |
|  | 3 | 100 | SRX703538 | 33 | 13.2 | 3,017,625 | 1,926,350 |
|  | 4 | 100 | SRX703508 | 33 | 13.2 | 3,131,836 | 1,817,067 |
|  | 5 | 100 | SRX703504 | 33 | 13.2 | 2,739,820 | 1,947,156 |
|  | 6 | 100 | SRX703511 | 33 | 13.2 | 2,997,191 | 1,956,779 |
|  | 7 | 100 | SRX703509 | 33 | 13.2 | 2,788,873 | 1,991,953 |
|  | 8 | 100 | SRX703532 | 33 | 13.2 | 2,853,844 | 1,923,391 |
|  | 9 | 100 | SRX703535 | 33 | 13.2 | 2,939,385 | 2,032,922 |
|  | 10 | 100 | SRX703537 | 33 | 13.2 | 3,079,185 | 1,879,770 |
|  | 11 | 100 | SRX703539 | 33 | 13.2 | 3,045,528 | 1,883,315 |
| L | 1 | 100 | SRX703541 | 32 | 12.8 | 3,054,680 | 3,089,734 |
|  | 2 | 100 | SRX703540 | 33 | 13.2 | 3,147,921 | 3,091,648 |
|  | 3 | 100 | SRX703542 | 33 | 13.2 | 3,074,258 | 3,135,548 |
|  | 4 | 100 | SRX703543 | 32 | 13.0 | 3,035,506 | 3,187,635 |
|  | 5 | 100 | SRX703548 | 33 | 13.2 | 3,553,960 | 2,825,913 |
|  | 6 | 100 | SRX703550 | 32.8 | 13.1 | 2,843,044 | 3,283,290 |
|  | 7 | 100 | SRX703552 | 33 | 13.2 | 3,818,360 | 2,727,436 |
|  | 8 | 100 | SRX703551 | 32.6 | 13.0 | 3,747,785 | 2,812,506 |
|  | 9 | 100 | SRX1544487 | 33 | 13.2 | 3,029,667 | 2,967,386 |
| Y | 1 | 100 | SRX703594 | 31.8 | 12.7 | 2,895,003 | 3,365,363 |
|  | 2 | 100 | SRX703596 | 33 | 13.2 | 3,646,253 | 2,814,432 |
|  | 3 | 100 | SRX703599 | 33 | 13.2 | 3,462,828 | 3,063,392 |
|  | 4 | 100 | SRX703597 | 33 | 13.2 | 3,511,229 | 3,076,198 |
|  | 5 | 100 | SRX703577 | 33 | 13.2 | 3,050,148 | 3,261,253 |
|  | 6 | 100 | SRX703579 | 33 | 13.2 | 3,210,946 | 3,054,866 |
|  | 7 | 100 | SRX703581 | 33 | 13.2 | 3,226,449 | 3,159,891 |
|  | 8 | 100 | SRX703582 | 33 | 13.2 | 3,331,057 | 3,026,034 |
|  | 9 | 100 | SRX703595 | 33 | 13.2 | 3,133,195 | 3,047,464 |
|  | 10 | 100 | SRX1544506 | 33 | 13.2 | 3,649,140 | 2,974,150 |
| average | - | - | - | 32.9 | 13.2 | - | - |
| Total | - | - | - | - | - | 12,564,044 | |

Table S2 Data summary and SNP statistics of the sequenced DLY pig family

| Breed / | Individual | High-quality | Genome | Q20 | Q30 | Genome coverage at least | | | Heterozygous | Homozygous |
| --- | --- | --- | --- | --- | --- | --- | --- | --- | --- | --- |
| Crossbred |  | data (Gb) | coverage (×) | (%) | (%) | 1 × (%) | 4 × (%) | 10 × (%) | SNP | SNP |
| D | 1 | 86.1 | 34.44 | 96.52 | 91.21 | 99.53 | 99.27 | 98.19 | 4,437,816 | 2,165,490 |
| L | 1 | 108.2 | 43.28 | 96.64 | 90.92 | 99.45 | 99.13 | 98.38 | 4,722,522 | 3,744,240 |
| Y | 1 | 90.4 | 36.16 | 95.58 | 89.22 | 99.16 | 98.85 | 98.01 | 5,444,996 | 3,399,276 |
| LY | 1 | 90.8 | 36.32 | 95.13 | 88.01 | 99.12 | 98.77 | 97.57 | 5,617,905 | 3,335,535 |
| DLY | 1 | 102.1 | 40.84 | 95.53 | 88.04 | 99.57 | 99.33 | 98.68 | 6,263,658 | 2,196,226 |
|  | 2 | 98.6 | 39.44 | 96.36 | 89.96 | 99.57 | 99.34 | 98.68 | 6,163,204 | 2,206,296 |
|  | 3 | 94.8 | 37.92 | 96.51 | 90.29 | 99.19 | 98.97 | 98.45 | 6,115,540 | 2,208,917 |
|  | 4 | 94.5 | 37.80 | 96.09 | 89.55 | 99.19 | 98.96 | 98.45 | 6,167,311 | 2,175,091 |
|  | 5 | 97.5 | 39.00 | 95.65 | 89.15 | 99.57 | 99.32 | 98.61 | 6,135,372 | 2,178,981 |
|  | 6 | 99.1 | 39.64 | 96.31 | 89.89 | 99.21 | 98.98 | 98.48 | 6,201,672 | 2,243,369 |
| Average | - | 96.2 | 38.48 | 96.03 | 89.62 | 99.36 | 99.09 | 98.35 | - | - |
| Total | - | 962 | - | - | - | - | - | - | 14,120,975 | |

Table S3 Validation of the accuracy of SNP identification by the chip

| **SNP type** | **Category** | **D** | **L** | **Y** | **LY** | **DLY** | | | | | |
| --- | --- | --- | --- | --- | --- | --- | --- | --- | --- | --- | --- |
|  |  |  |  |  |  | **1** | **2** | **3** | **4** | **5** | **6** |
| Homozygous | SNPs identified by the chip | 4,323 | 8,740 | 7,560 | 7,698 | 4,586 | 4,742 | 4,464 | 4,447 | 4,645 | 4,549 |
|  | Validated by GATK | 4,210 | 8,591 | 7,429 | 7,584 | 4,485 | 4,655 | 4,390 | 4,365 | 4,553 | 4,465 |
|  | Concordance ratio (%) | 97.39 | 98.30 | 98.27 | 98.52 | 97.80 | 98.17 | 98.34 | 98.16 | 98.02 | 98.15 |
| Heterozygous | SNPs identified by the chip | 11,422 | 11,552 | 14,223 | 13,870 | 15,843 | 15,484 | 15,768 | 15,766 | 15,610 | 16,033 |
|  | Validated by GATK | 11,254 | 11,303 | 13,873 | 13,664 | 15,613 | 15,243 | 15,527 | 15,528 | 15,390 | 15,814 |
|  | Concordance ratio (%) | 98.53 | 97.84 | 97.53 | 98.51 | 98.55 | 98.44 | 98.47 | 98.49 | 98.59 | 98.63 |
| Total | SNPs identified by the chip | 15,745 | 20,292 | 21,783 | 21,568 | 20429 | 20,226 | 20,232 | 20,213 | 20,255 | 20,582 |
|  | Validated by GATK | 15,464 | 19,894 | 21,300 | 21,248 | 20098 | 19,898 | 19,917 | 19,893 | 19,943 | 20,279 |
|  | Concordance ratio (%) | 98.22 | 98.04 | 97.78 | 98.52 | 98.38 | 98.38 | 98.44 | 98.42 | 98.46 | 98.53 |

Table S4 Statistics of occurrence for five different genomic elements

| **Category** | | **Total detected SNPs of three pig breeds** | | | **SNPs of breed-of-origin of alleles** | | |
| --- | --- | --- | --- | --- | --- | --- | --- |
|  |  | **Number** | **Occurrence (%)** | **Gene** | **Number** | **Occurrence (%)** | **Gene** |
| Total | | 12,564,687 | 100 | - | 529,930 | 100 | - |
| Upstream | | 92,083 | 0.73 | 14,646 | 3,918 | 0.74 | 1,380 |
| Downstream | | 194,780 | 1.55 | 17,102 | 8,544 | 1.61 | 1,993 |
| Intergenic | | 7,281,054 | 57.95 | 14,814 | 299,050 | 56.43 | 5,426 |
| Intronic | | 4,921,514 | 39.17 | 17,740 | 214,952 | 40.56 | 5,602 |
| Exonic | Nonsynonymous | 26,592 | 0.21 | 8,761 | 1,165 | 0.22 | 664 |
|  | Stop gain | 332 | ~0 | 309 | 12 | ~0 | 12 |
|  | Stop loss | 75 | ~0 | 74 | 5 | ~0 | 4 |
|  | Synonymous | 43,453 | 0.35 | 11,785 | 1,983 | 0.37 | 1,036 |

Table S5 Function enrichment (GO and KEGG) for breed-of-origin of alleles located in upstream region

| **Function Category** | **Count** | ***P*-value** | **Fold Enrichment** |
| --- | --- | --- | --- |
| GO:0005758 mitochondrial intermembrane space | 13 | 6.1 x 10^-4^ | 3.23 |
| GO:0005515 protein binding | 535 | 1.3 x 10^-3^ | 1.09 |
| GO:0000228 nuclear chromosome | 10 | 2.1 x 10^-3^ | 3.47 |
| GO:0001594 trace-amine receptor activity | 4 | 3.1 x 10^-3^ | 11.97 |
| GO:0070062 extracellular exosome | 185 | 3.3 x 10^-3^ | 1.21 |
| GO:0005739 mitochondrion | 96 | 3.7 x 10^-3^ | 1.33 |
| GO:0097110 scaffold protein binding | 9 | 4.6 x 10^-3^ | 3.37 |
| GO:0046329 negative regulation of JNK cascade | 6 | 6.1 x 10^-3^ | 4.93 |
| BIOCARTA Stat3 Signaling Pathway | 4 | 6.4 x 10^-3^ | 9.45 |
| GO:0016042 lipid catabolic process | 12 | 6.9 x 10^-3^ | 2.55 |
| GO:0007217 tachykinin receptor signaling pathway | 4 | 7.6 x 10^-3^ | 9.05 |
| GO:0005829 cytosol | 210 | 9.3 x 10^-3^ | 1.16 |
| GO:0006470 protein dephosphorylation | 15 | 9.7 x 10^-3^ | 2.15 |
| GO:0030176 integral component of endoplasmic reticulum membrane | 13 | 0.011 | 2.29 |
| GO:0005198 structural molecule activity | 24 | 0.011 | 1.75 |
| GO:0046007 negative regulation of activated T cell proliferation | 4 | 0.011 | 8.04 |
| GO:0007059 chromosome segregation | 10 | 0.012 | 2.66 |
| GO:0043410 positive regulation of MAPK cascade | 11 | 0.014 | 2.46 |
| GO:0019953 sexual reproduction | 4 | 0.015 | 7.24 |
| GO:0004177 aminopeptidase activity | 6 | 0.015 | 3.99 |
| GO:0008190 eukaryotic initiation factor 4E binding | 4 | 0.015 | 7.18 |
| GO:0055085 transmembrane transport | 23 | 0.016 | 1.71 |
| GO:0019784 NEDD8-specific protease activity | 3 | 0.017 | 13.46 |
| GO:0004527 exonuclease activity | 5 | 0.019 | 4.73 |
| GO:0045815 positive regulation of gene expression, epigenetic | 9 | 0.021 | 2.63 |
| GO:0016290 palmitoyl-CoA hydrolase activity | 4 | 0.021 | 6.53 |
| GO:0038083 peptidyl-tyrosine autophosphorylation | 7 | 0.021 | 3.17 |
| GO:0007155 cell adhesion | 37 | 0.022 | 1.46 |
| GO:0003723 RNA binding | 43 | 0.022 | 1.41 |
| GO:0031901 early endosome membrane | 13 | 0.024 | 2.06 |
| GO:0016032 viral process | 26 | 0.026 | 1.57 |
| GO:0016887 ATPase activity | 18 | 0.027 | 1.77 |
| GO:0004298 threonine-type endopeptidase activity | 5 | 0.027 | 4.28 |
| GO:0051290 protein heterotetramerization | 7 | 0.027 | 3.02 |
| GO:0045039 protein import into mitochondrial inner membrane | 3 | 0.027 | 10.86 |
| GO:0010035 response to inorganic substance | 3 | 0.027 | 10.86 |
| GO:0071013 catalytic step 2 spliceosome | 11 | 0.028 | 2.19 |
| GO:0030864 cortical actin cytoskeleton | 7 | 0.028 | 2.99 |
| GO:0046677 response to antibiotic | 6 | 0.029 | 3.39 |
| GO:0007601 visual perception | 19 | 0.029 | 1.71 |
| GO:0033209 tumor necrosis factor-mediated signaling pathway | 13 | 0.029 | 1.99 |
| GO:0050901 leukocyte tethering or rolling | 4 | 0.032 | 5.57 |
| GO:0097284 hepatocyte apoptotic process | 4 | 0.032 | 5.57 |
| GO:0045121 membrane raft | 19 | 0.032 | 1.69 |
| GO:0005856 cytoskeleton | 30 | 0.033 | 1.49 |
| GO:0007173 epidermal growth factor receptor signaling pathway | 8 | 0.034 | 2.58 |
| GO:0004872 receptor activity | 20 | 0.034 | 1.66 |
| GO:0043005 neuron projection | 21 | 0.034 | 1.63 |
| GO:0031593 polyubiquitin binding | 5 | 0.036 | 3.91 |
| GO:0005925 focal adhesion | 31 | 0.037 | 1.46 |
| GO:0020027 hemoglobin metabolic process | 3 | 0.039 | 9.05 |
| GO:0046907 intracellular transport | 5 | 0.041 | 3.77 |
| GO:0032456 endocytic recycling | 5 | 0.041 | 3.77 |
| GO:0006508 proteolysis | 38 | 0.042 | 1.38 |
| GO:0031047 gene silencing by RNA | 12 | 0.043 | 1.96 |
| GO:0005882 intermediate filament | 12 | 0.044 | 1.95 |
| GO:0006352 DNA-templated transcription, initiation | 6 | 0.046 | 3.02 |
| GO:0006112 energy reserve metabolic process | 4 | 0.046 | 4.83 |
| GO:0097009 energy homeostasis | 4 | 0.046 | 4.83 |
| GO:0047617 acyl-CoA hydrolase activity | 4 | 0.047 | 4.79 |
| GO:0051603 proteolysis involved in cellular protein catabolic process | 7 | 0.047 | 2.64 |
| GO:0016310 phosphorylation | 11 | 0.049 | 1.99 |

Table S6 Function enrichment (GO and KEGG) for breed-of-origin of alleles located in downstream region

| **Function Category** | **Count** | ***P*-value** | **Fold Enrichment** |
| --- | --- | --- | --- |
| GO:0005515 protein binding | 824 | 1.7 x 10^-4^ | 1.09 |
| GO:0012507 ER to Golgi transport vesicle membrane | 14 | 2.9 x 10^-4^ | 3.19 |
| GO:0032395 MHC class II receptor activity | 7 | 1.1 x 10^-3^ | 5.41 |
| GO:0060687 regulation of branching involved in prostate gland morphogenesis | 5 | 1.6 x 10^-3^ | 8.27 |
| GO:0006470 protein dephosphorylation | 22 | 2.6 x 10^-3^ | 2.02 |
| GO:0043005 neuron projection | 34 | 3.1 x 10^-3^ | 1.69 |
| GO:0009966 regulation of signal transduction | 9 | 3.2 x 10^-3^ | 3.47 |
| GO:0042803 protein homodimerization activity | 85 | 3.6 x 10^-3^ | 1.35 |
| GO:0050790 regulation of catalytic activity | 14 | 3.8 x 10^-3^ | 2.45 |
| GO:0006886 intracellular protein transport | 34 | 3.9 x 10^-3^ | 1.67 |
| GO:0051216 cartilage development | 13 | 4.1 x 10^-3^ | 2.55 |
| GO:0048705 skeletal system morphogenesis | 9 | 4.9 x 10^-3^ | 3.25 |
| GO:0031175 neuron projection development | 18 | 5.1 x 10^-3^ | 2.08 |
| GO:0016192 vesicle-mediated transport | 24 | 5.7 x 10^-3^ | 1.83 |
| GO:0060333 interferon-gamma-mediated signaling pathway | 14 | 7.2 x 10^-3^ | 2.28 |
| GO:1900006 positive regulation of dendrite development | 6 | 9.3 x 10^-3^ | 4.34 |
| GO:0061028 establishment of endothelial barrier | 6 | 9.3 x 10^-3^ | 4.34 |
| GO:0048008 platelet-derived growth factor receptor signaling pathway | 8 | 0.011 | 3.19 |
| GO:0006869 lipid transport | 14 | 0.013 | 2.13 |
| GO:0071541 eukaryotic translation initiation factor 3 complex, eIF3m | 4 | 0.016 | 6.76 |
| GO:0007173 epidermal growth factor receptor signaling pathway | 11 | 0.021 | 2.27 |
| GO:0021502 neural fold elevation formation | 3 | 0.021 | 11.57 |
| GO:0032526 response to retinoic acid | 9 | 0.022 | 2.54 |
| GO:0051592 response to calcium ion | 11 | 0.026 | 2.19 |
| GO:0010518 positive regulation of phospholipase activity | 4 | 0.026 | 5.79 |
| GO:0007626 locomotory behavior | 14 | 0.028 | 1.93 |
| GO:0001750 photoreceptor outer segment | 10 | 0.029 | 2.28 |
| GO:0042127 regulation of cell proliferation | 25 | 0.029 | 1.56 |
| GO:0003281 ventricular septum development | 7 | 0.029 | 2.89 |
| GO:0006198 cAMP catabolic process | 5 | 0.035 | 3.86 |
| GO:0070700 BMP receptor binding | 4 | 0.036 | 5.15 |
| GO:0034605 cellular response to heat | 8 | 0.037 | 2.51 |
| GO:0003170 heart valve development | 3 | 0.039 | 8.68 |
| GO:0048619 embryonic hindgut morphogenesis | 3 | 0.039 | 8.68 |
| GO:0071260 cellular response to mechanical stimulus | 12 | 0.041 | 1.96 |
| GO:0019933 cAMP-mediated signaling | 8 | 0.042 | 2.44 |
| GO:0000398 mRNA splicing, via spliceosome | 28 | 0.044 | 1.46 |
| KEGG Neuroactive ligand-receptor interaction | 36 | 0.045 | 1.37 |
| GO:0006814 sodium ion transport | 13 | 0.045 | 1.86 |
| GO:0005758 mitochondrial intermembrane space | 12 | 0.046 | 1.92 |
| GO:0005916 fascia adherens | 4 | 0.046 | 4.73 |
| GO:0030032 lamellipodium assembly | 7 | 0.047 | 2.61 |
| GO:0008277 regulation of G-protein coupled receptor protein signaling pathway | 8 | 0.047 | 2.38 |
| GO:0008013 beta-catenin binding | 13 | 0.048 | 1.84 |
| GO:0009791 post-embryonic development | 12 | 0.048 | 1.91 |
| GO:0060174 limb bud formation | 4 | 0.049 | 4.63 |
| GO:1903206 negative regulation of hydrogen peroxide-induced cell death | 4 | 0.049 | 4.63 |
| GO:0051901 positive regulation of mitochondrial depolarization | 4 | 0.049 | 4.63 |
| GO:0046415 urate metabolic process | 4 | 0.049 | 4.63 |
| GO:0035646 endosome to melanosome transport | 4 | 0.049 | 4.63 |
| GO:0042493 response to drug | 36 | 0.049 | 1.37 |
| GO:0008380 RNA splicing | 22 | 0.049 | 1.53 |
| GO:0050680 negative regulation of epithelial cell proliferation | 10 | 0.049 | 2.07 |
| GO:0045177 apical part of cell | 12 | 0.049 | 1.89 |
| GO:0031143 pseudopodium | 5 | 0.049 | 3.48 |

Table S7 Function enrichment (GO and KEGG) for breed-of-origin of alleles located in exonic region

| **Function Category** | **Count** | ***P*-value** | **Fold Enrichment** |
| --- | --- | --- | --- |
| GO:0005604 basement membrane | 18 | 3.1 x 10^-6^ | 3.82 |
| GO:0016887 ATPase activity | 29 | 4.9 x 10^-6^ | 2.63 |
| GO:0000242 pericentriolar material | 8 | 6.9 x 10^-5^ | 7.07 |
| GO:0005578 proteinaceous extracellular matrix | 33 | 1.4 x 10^-4^ | 2.07 |
| GO:0005085 guanyl-nucleotide exchange factor activity | 18 | 7.1 x 10^-4^ | 2.53 |
| GO:0012507 ER to Golgi transport vesicle membrane | 11 | 8.6 x 10^-4^ | 3.55 |
| GO:0005509 calcium ion binding | 65 | 1.1 x 10^-3^ | 1.51 |
| GO:0032395 MHC class II receptor activity | 6 | 1.4 x 10^-3^ | 6.63 |
| GO:0005539 glycosaminoglycan binding | 6 | 3.5 x 10^-3^ | 5.52 |
| GO:0019886 antigen processing and presentation of exogenous peptide antigen via MHC class II | 14 | 3.8 x 10^-3^ | 2.49 |
| GO:0016324 apical plasma membrane | 30 | 4.7 x 10^-3^ | 1.73 |
| GO:0045121 membrane raft | 23 | 5.7 x 10^-3^ | 1.87 |
| GO:0005201 extracellular matrix structural constituent | 11 | 6.6 x 10^-3^ | 2.72 |
| GO:0045860 positive regulation of protein kinase activity | 9 | 7.1 x 10^-3^ | 3.13 |
| GO:0043005 neuron projection | 25 | 7.8 x 10^-3^ | 1.77 |
| GO:0004222 metalloendopeptidase activity | 15 | 8.1 x 10^-3^ | 2.19 |
| GO:0042613 MHC class II protein complex | 6 | 8.3 x 10^-3^ | 4.57 |
| GO:0016459 myosin complex | 9 | 8.9 x 10^-3^ | 3.02 |
| GO:0045177 apical part of cell | 11 | 0.013 | 2.46 |
| GO:0055085 transmembrane transport | 25 | 0.014 | 1.68 |
| GO:0005198 structural molecule activity | 25 | 0.014 | 1.68 |
| KEGG Neuroactive ligand-receptor interaction | 29 | 0.015 | 1.58 |
| GO:0016311 dephosphorylation | 12 | 0.015 | 2.29 |
| GO:0002504 antigen processing and presentation of peptide or polysaccharide antigen via MHC class II | 5 | 0.017 | 4.81 |
| KEGG Pancreatic secretion | 13 | 0.019 | 2.11 |
| GO:0005516 calmodulin binding | 20 | 0.019 | 1.75 |
| GO:0006970 response to osmotic stress | 5 | 0.021 | 4.55 |
| GO:0007616 long-term memory | 6 | 0.026 | 3.51 |
| GO:0014069 postsynaptic density | 19 | 0.026 | 1.73 |
| KEGG Antigen processing and presentation | 11 | 0.027 | 2.19 |
| GO:0060333 interferon-gamma-mediated signaling pathway | 10 | 0.028 | 2.31 |
| BIOCARTA Antigen Processing and Presentation | 4 | 0.029 | 5.63 |
| GO:0048008 platelet-derived growth factor receptor signaling pathway | 6 | 0.029 | 3.39 |
| GO:0048208 COPII vesicle coating | 9 | 0.031 | 2.41 |
| GO:0030669 clathrin-coated endocytic vesicle membrane | 7 | 0.033 | 2.87 |
| GO:0045111 intermediate filament cytoskeleton | 8 | 0.034 | 2.58 |
| GO:0016042 lipid catabolic process | 11 | 0.034 | 2.12 |
| GO:0050896 response to stimulus | 9 | 0.034 | 2.38 |
| GO:0005871 kinesin complex | 8 | 0.037 | 2.53 |
| GO:0016055 Wnt signaling pathway | 19 | 0.037 | 1.66 |
| KEGG Adipocytokine signaling pathway | 10 | 0.039 | 2.16 |
| KEGG Thyroid hormone synthesis | 10 | 0.039 | 2.16 |
| GO:0045598 regulation of fat cell differentiation | 4 | 0.041 | 5.04 |
| GO:0045202 synapse | 18 | 0.042 | 1.67 |
| GO:0046677 response to antibiotic | 6 | 0.043 | 3.07 |
| GO:0048705 skeletal system morphogenesis | 6 | 0.043 | 3.07 |
| GO:0046330 positive regulation of JNK cascade | 9 | 0.043 | 2.27 |
| GO:0006470 protein dephosphorylation | 14 | 0.044 | 1.82 |
| GO:0030425 dendrite | 29 | 0.045 | 1.45 |
| GO:0006644 phospholipid metabolic process | 8 | 0.045 | 2.42 |
| GO:0000784 nuclear chromosome, telomeric region | 14 | 0.046 | 1.81 |
| GO:0005581 collagen trimer | 11 | 0.047 | 2.01 |
| GO:0042626 ATPase activity, coupled to transmembrane movement of substances | 7 | 0.047 | 2.64 |
| GO:0031295 T cell costimulation | 10 | 0.048 | 2.09 |
| GO:0046835 carbohydrate phosphorylation | 5 | 0.049 | 3.56 |

Table S8 Function enrichment (GO and KEGG) for breed-of-origin of alleles located in intronic region

| **Function Category** | **Count** | ***P*-value** | **Fold Enrichment** |
| --- | --- | --- | --- |
| KEGG Glutamatergic synapse | 22 | 3.1 x 10^-7^ | 3.71 |
| GO:0005509 calcium ion binding | 62 | 1.1 x 10^-5^ | 1.79 |
| KEGG Gastric acid secretion | 15 | 1.8 x 10^-5^ | 3.94 |
| KEGG Adrenergic signaling in cardiomyocytes | 22 | 1.8 x 10^-5^ | 2.89 |
| KEGG Insulin secretion | 16 | 2.6 x 10^-5^ | 3.61 |
| KEGG Dopaminergic synapse | 20 | 2.9 x 10^-5^ | 2.99 |
| KEGG Thyroid hormone synthesis | 14 | 5.2 x 10^-5^ | 3.84 |
| KEGG GnRH signaling pathway | 16 | 5.9 x 10^-5^ | 3.37 |
| GO:0030032 lamellipodium assembly | 9 | 9.6 x 10^-5^ | 5.89 |
| KEGG Cholinergic synapse | 17 | 1.9 x 10^-4^ | 2.94 |
| KEGG Aldosterone synthesis and secretion | 14 | 2.4 x 10^-4^ | 3.32 |
| GO:0043547 positive regulation of GTPase activity | 48 | 3.2 x 10^-4^ | 1.72 |
| KEGG Vascular smooth muscle contraction | 17 | 4.1 x 10^-4^ | 2.74 |
| GO:0051592 response to calcium ion | 11 | 4.8 x 10^-4^ | 3.84 |
| KEGG Inflammatory mediator regulation of TRP channels | 15 | 4.9 x 10^-4^ | 2.94 |
| KEGG Oxytocin signaling pathway | 20 | 5.1 x 10^-4^ | 2.43 |
| GO:0016477 cell migration | 20 | 8.4 x 10^-4^ | 2.36 |
| KEGG Long-term depression | 11 | 9.3 x 10^-4^ | 3.52 |
| GO:0007229 integrin-mediated signaling pathway | 14 | 1.1 x 10^-3^ | 2.87 |
| GO:0005539 glycosaminoglycan binding | 6 | 1.3 x 10^-3^ | 6.92 |
| GO:0048471 perinuclear region of cytoplasm | 47 | 1.8 x 10^-3^ | 1.59 |
| GO:0030036 actin cytoskeleton organization | 16 | 1.8 x 10^-3^ | 2.49 |
| GO:0004222 metalloendopeptidase activity | 14 | 3.1 x 10^-3^ | 2.57 |
| GO:0038083 peptidyl-tyrosine autophosphorylation | 8 | 3.1 x 10^-3^ | 4.06 |
| KEGG Calcium signaling pathway | 19 | 5.3 x 10^-3^ | 2.04 |
| KEGG Melanogenesis | 13 | 5.4 x 10^-3^ | 2.49 |
| GO:0086091 regulation of heart rate by cardiac conduction | 7 | 6.7 x 10^-3^ | 4.06 |
| KEGG Bile secretion | 10 | 8.9 x 10^-3^ | 2.78 |
| GO:0000287 magnesium ion binding | 19 | 9.7 x 10^-3^ | 1.93 |
| GO:0042127 regulation of cell proliferation | 18 | 0.011 | 1.97 |
| GO:0019933 cAMP-mediated signaling | 7 | 0.011 | 3.74 |
| GO:0010811 positive regulation of cell-substrate adhesion | 7 | 0.011 | 3.74 |
| GO:0072659 protein localization to plasma membrane | 9 | 0.012 | 2.89 |
| KEGG Ovarian steroidogenesis | 8 | 0.013 | 3.13 |
| GO:0008217 regulation of blood pressure | 9 | 0.014 | 2.81 |
| KEGG ECM-receptor interaction | 11 | 0.014 | 2.43 |
| GO:0007268 chemical synaptic transmission | 21 | 0.015 | 1.77 |
| GO:0030667 secretory granule membrane | 5 | 0.016 | 5.02 |
| GO:0005783 endoplasmic reticulum | 54 | 0.016 | 1.38 |
| GO:0035338 long-chain fatty-acyl-CoA biosynthetic process | 7 | 0.016 | 3.38 |
| GO:0050796 regulation of insulin secretion | 9 | 0.017 | 2.72 |
| KEGG Sphingolipid signaling pathway | 13 | 0.022 | 2.08 |
| GO:0030426 growth cone | 12 | 0.022 | 2.18 |
| GO:0035335 peptidyl-tyrosine dephosphorylation | 11 | 0.024 | 2.26 |
| GO:0006886 intracellular protein transport | 20 | 0.025 | 1.72 |
| KEGG Regulation of lipolysis in adipocytes | 8 | 0.025 | 2.74 |
| GO:0031175 neuron projection development | 11 | 0.025 | 2.23 |
| KEGG Oocyte meiosis | 12 | 0.026 | 2.11 |
| KEGG Regulation of actin cytoskeleton | 19 | 0.026 | 1.73 |
| KEGG cAMP signaling pathway | 18 | 0.028 | 1.74 |
| KEGG Phosphatidylinositol signaling system | 11 | 0.031 | 2.05 |
| GO:0007605 sensory perception of sound | 13 | 0.031 | 1.98 |
| GO:0050896 response to stimulus | 8 | 0.032 | 2.62 |
| KEGG Estrogen signaling pathway | 11 | 0.032 | 2.13 |
| KEGG Glucagon signaling pathway | 11 | 0.032 | 2.13 |
| GO:0004871 signal transducer activity | 17 | 0.038 | 1.73 |
| GO:0003824 catalytic activity | 16 | 0.038 | 1.77 |
| GO:0008277 regulation of G-protein coupled receptor protein signaling pathway | 6 | 0.041 | 3.12 |
| KEGG Rap1 signaling pathway | 18 | 0.046 | 1.65 |
| GO:0007156 homophilic cell adhesion via plasma membrane adhesion molecules | 14 | 0.048 | 1.79 |
| GO:0031901 early endosome membrane | 11 | 0.049 | 2.01 |

Table S9 Summary of the 50 genes containing nonsynonymous SNPs of breed-of-origin of alleles with highest probability of heterozygous SNP

| **Gene Symbol** | **Gene name** | **Chromosome** | **Position** | **Probability of**  **heterozygous SNP** |
| --- | --- | --- | --- | --- |
| *AKAP9* | A-kinase anchoring protein 9 | 9 | 72,027,468 | 0.98 |
| *ENDOU* | Endonuclease, poly(U) specific | 5 | 78,083,854 | 0.94 |
| *FAM221A* | Family with sequence similarity 221 | 18 | 48,393,269 | 0.9 |
| *AHI1* | Abelson helper integration site 1 | 1 | 28,589,827 | 0.9 |
| *USP20* | Ubiquitin specific peptidase 20 | 1 | 270,017,455 | 0.9 |
| *LYZ* | Lysozyme | 5 | 33,617,134 | 0.89 |
| *MCIDAS* | Multiciliate differentiation and DNA synthesis associated cell cycle protein | 16 | 34,383,925 | 0.89 |
| *ASXL3* | ASXL transcriptional regulator 3 | 6 | 117,515,919 | 0.88 |
| *BORA* | Bora, aurora kinase A activator | 11 | 45,032,978 | 0.88 |
| *METAP1D* | Methionyl aminopeptidase type 1D | 15 | 78,181,771 | 0.86 |
| *OCA2* | Oculocutaneous albinism II | 15 | 56,768,830 | 0.86 |
| *MARVELD3* | MARVEL domain containing 3 | 6 | 14,629,824 | 0.86 |
| *CORIN* | Corin, serine peptidase | 8 | 37,545,419 | 0.86 |
| *IFNLR1* | Interferon lambda receptor 1 | 6 | 81,902,375 | 0.86 |
| *DNAJB12* | DnaJ heat shock protein family (Hsp40) member B12 | 14 | 75,180,371 | 0.86 |
| *ADAMTS18* | ADAM metallopeptidase with thrombospondin type 1 motif 18 | 6 | 10,623,104 | 0.85 |
| *CEP57L1* | Centrosomal protein 57 like 1 | 1 | 75,223,360 | 0.84 |
| *EPHA10* | EPH receptor A10 | 6 | 93,800,268 | 0.84 |
| *TCHHL1* | Trichohyalin like 1 | 4 | 97,179,656 | 0.84 |
| *POGZ* | Pogo transposable element derived with ZNF domain | 4 | 97,829,637 | 0.84 |
| *STK31* | Serine/threonine kinase 31 | 18 | 48,342,617 | 0.84 |
| *NUDT13* | Nudix hydrolase 13 | 14 | 75,945,482 | 0.84 |
| *NVL* | Nuclear VCP-like | 10 | 12,661,964 | 0.84 |
| *NEU2* | Neuraminidase 2 | 15 | 133,475,330 | 0.84 |
| *UBE3C* | Ubiquitin protein ligase E3C | 18 | 1,630,010 | 0.83 |
| *TRPM6* | Transient receptor potential cation channel subfamily M member 6 | 1 | 227,843,764 | 0.83 |
| *CLYBL* | Citrate lyase beta like | 11 | 68,738,893 | 0.83 |
| *DUPD1* | Dual specificity phosphatase and pro isomerase domain containing 1 | 14 | 77,603,362 | 0.83 |
| *MAP2* | Microtubule-associated protein 2 | 15 | 112,423,394 | 0.83 |
| *CRIM1* | Cysteine rich transmembrane BMP regulator 1 | 3 | 103,773,071 | 0.82 |
| *ZSWIM8* | Zinc finger SWIM-type containing 8 | 14 | 76,521,336 | 0.82 |
| *LAMC3* | Laminin gamma 3 | 1 | 271,045,921 | 0.82 |
| *TNXB* | Tenascin XB | 7 | 24,150,524 | 0.82 |
| *GJB5* | Gap junction beta-5 | 6 | 91,002,967 | 0.82 |
| *ANAPC1* | Anaphase promoting complex subunit 1 | 3 | 44,586,240 | 0.82 |
| *NVL* | Nuclear VCP-like | 10 | 12,659,643 | 0.82 |
| *USP40* | Ubiquitin specific peptidase 40 | 15 | 133,890,924 | 0.81 |
| *CYB5R4* | Cytochrome b5 reductase 4 | 1 | 53,190,625 | 0.81 |
| *HERC6* | HECT and RLD domain containing E3 ubiquitin protein ligase family member 6 | 8 | 130,624,681 | 0.81 |
| *LINGO4* | Leucine rich repeat and Ig domain containing 4 | 4 | 97,404,380 | 0.81 |
| *SLIT3* | Slit guidance ligand 3 | 16 | 55,214,924 | 0.81 |
| *ZFHX3* | Zinc finger homeobox 3 | 6 | 15,861,722 | 0.8 |
| *SIMC1* | SUMO interacting motifs containing 1 | 2 | 81,540,468 | 0.8 |
| *CRNN* | Cornulin | 4 | 96,935,328 | 0.8 |
| *ARHGEF28* | Rho guanine nucleotide exchange factor 28 | 2 | 82,986,820 | 0.8 |
| *HARS2* | Histidyl-tRNA synthetase 2 | 2 | 142,405,520 | 0.8 |
| *ASCC1* | Activating signal cointegrator 1 | 14 | 75,080,922 | 0.8 |
| *ACSF2* | Acyl-CoA synthetase family member 2 | 12 | 26,739,774 | 0.8 |
| *CENPE* | Centromere protein E | 8 | 117,908,657 | 0.8 |
| *MFSD2A* | Major facilitator superfamily domain containing 2A | 6 | 95,748,552 | 0.8 |

Table S10 Summary of transcriptome data of the sequenced 4 tissues

| **DLY** | **Adipose** | | **Skeletal muscle** | | **Liver** | | **Heart** | |
| --- | --- | --- | --- | --- | --- | --- | --- | --- |
|  | High-quality  data (Gb) | Mapping  rate (%) | High-quality  data (Gb) | Mapping  rate (%) | High-quality  data (Gb) | Mapping  rate (%) | High-quality  data (Gb) | Mapping  rate (%) |
| 1 | 5.9 | 81.5 | 3.6 | 84.7 | 4.8 | 82.3 | 5.1 | 81.3 |
| 2 | 4.4 | 81.7 | 3.8 | 84.5 | 4.7 | 82.8 | 4.7 | 81.5 |
| 3 | 4.7 | 80.0 | 4.7 | 80.0 | 4.8 | 83.4 | 4.3 | 81.0 |
| 4 | 4.7 | 86.2 | 4.2 | 85.9 | 4.6 | 87.7 | 4.6 | 86.0 |
| 5 | 4.3 | 86.5 | 5.1 | 85.8 | 4.9 | 87.0 | 4.4 | 86.0 |
| 6 | 5.1 | 86.3 | 4.9 | 85.4 | 5.1 | 87.6 | 4.1 | 86.1 |
